# Autonomous Learning from Intrinsic Sensory Dependencies Yields Generalizable Representations in Cortical-like Networks

**DOI:** 10.1101/2024.07.17.603881

**Authors:** Udaya B. Rongala, Henrik Jörntell

**Affiliations:** Neural Basis of Sensorimotor Control, Department of Experimental Medical Science, Lund University, Lund, Sweden

## Abstract

How biological brains become operational from development and onwards is an unresolved issue. Here we explored the emerging effects in a brain circuit model that received simulated touches through biological skin model, which featured the potentially critical aspect of sensory dependencies resulting from mechanical coupling across the tissue. Our skin model system was connected to a recurrent network a generic, primitive cortical-like subnetwork that was composed of fully connected excitatory and inhibitory neurons, where each individual synapse was subject to continuous, independent, Hebbian-like learning. We used continuous random mechanical activations of the skin model, in this regard mimicking behavioral patterns of early brain development observed in infants, and let the neuron-independent learning define the function that emerged in the network. Remarkably, we found that the network could rapidly learn to separate various naive dynamic skin inputs and solve a kinematics task it had never encountered, even when substantial parts of the sensor population or even network connections were removed post-training. We propose that autonomous learning from a sensor population with intrinsic dependencies could cause the extensively recursive cortical network to gradually adapt its intrinsic dynamics to better mirror the various dynamics of whatever body it is connected to, which result in many biologically useful features for the early acquisition of brain circuitry function.

## Introduction

A striking difference between biology and artificial intelligence is that the brain seems to be learning effectively and in a generalizable manner from fewer examples and less repetitious inputs (1). Why that happens is an unresolved issue, but could depend on a range of issues such as learning strategies (2), what specific network parameters that the learning is altering and more. One factor that has been largely overlooked is that the brain learns about the world through the lens of the biomechanical properties of its body. The body is composed of tissues with dynamic compliance, such as in the skin (3) but also in deep tissues in general, which make the whole body mechanically coupled (4). Because the sensors are distributed within such compliant tissues they can display complex patterns of dependencies, which can be demonstrated by the time-varying entropy in the structure of the sensor population signal (3). The mechanical couplings will also result in various spatiotemporal regularities in the sensor population signals. Combining information expressed on the bases of known regularities can be used to describe also previously unexperienced real-world interactions by means of the principle of superposition (3, 5, 6, 7, 8). A network learning strategy that focuses on representing (i.e. to symbolize or to provide a quantitative measure of) such regularities as broadly as possible could therefore become more generalizable (7) and more robust to sensor loss and/or sensor drift as well as localized brain damage, all of which would be valuable features in biological life and evolution.

How a cortical-like network would evolve its functionality through learning when connected to a biomechanical system with such sensory dependencies is not known. An intersting aspect is that the cortical network differs from other parts of the central nervous system by having more extensive recursive connectivity (in contrast, the networks in the brainstem nuclei and the cerebellum are more feed-forward in character (8, 9)). This can be of an advantage to more fully capture complex sensory dependencies, where mass-induced inertial effects across both body and local tissues introduce time-dependent factors to those dependencies (3, 4). Both the machine learning and the neurocomputation literature are full of examples of different types of recurrent neural networks (RNNs). In fact, networks with extensive recurrent connections, similar to Reservoir Computing (RC) or Echo State (ESN) networks, have been taken as reflective of the general principles of cortical network operation (10, 11). However, training of RC and ESN networks have almost exclusively been applied to the incoming or the outgoing connections from that network, and not within the recursive network itself, whereas RNN training typically uses external supervision, backpropagation or similar non-intrinsically driven learning. Network-intrinsic learning has sometimes been implemented in the neurocomputation literature, but then instead typically under the condition of isolating the excitatory or the inhibitory synaptic plasticity during the learning process or considering inhibitory synapse learning as a homeostatic reaction to the learning in the excitatory synapses (12, 13, 14, 15, 16, 17, 18).

In this paper, rather than imposing any *a priori* external goals for the learning, we wanted to study the emerging effects of autonomous learning in a fully connected network, exposed to random activations of a sensory system with extensive intrinsic mechanical couplings. This Hebbian-like learning was unrestrained and ran in parallel across all excitatory and inhibitory synapses. We wanted to do this using a previously published neuron model that includes an emulation of the membrane time constant of biological neurons, which we have previously shown can eliminate potential sources of instability in recurrent networks (19), which can be important for correlation-based (i.e. Hebbian-like) learning to not amplify instability. Emerging effects are important as they may better reflect the complex operation of brain circuitry *in vivo*, since the brain, as a complex entity, exhibits intrinsic properties and behaviors that its individual parts do not possess on their own — properties that can only emerge when these parts interact according to their own local rules. Our aim was to study these emergent effects in the simplest possible network that would still be capturing central functional principles compatible with a primitive cortical circuitry, i.e. early in development or early in evolution (20, 19, 21). Similarly, we were more interested in circuitry features conveying overarching functional effects rather than *a priori* copying the histological structure of the cortex. The biological motivations for why our network architecture (Fig. 1A) can be considered cortical-like is provided in the Methods.

**Figure 1:**
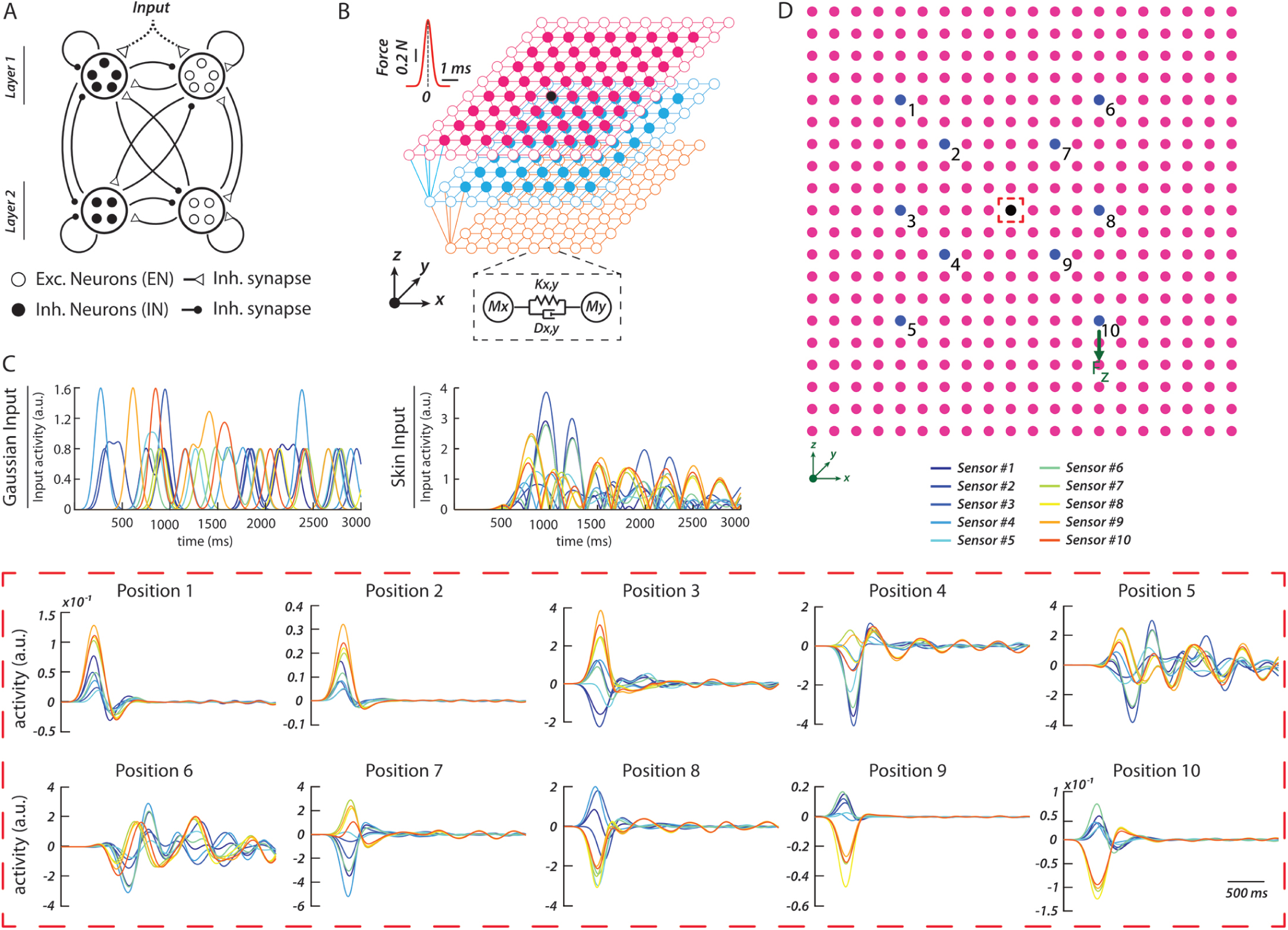
Network structure and training inputs from the dynamic skin tissue model. (**A**) Schematic of the fully connected network structure. (**B**) Three-layer damped mass-spring skin model (3) used to generate the intrinsically dependent sensory inputs. Top left corner, time profile of the applied forces for the different inputs. Masses, shown as nodes, were connected by springs and dampers represented as links (internode distance 1.0 mm). Note that the actual model was larger and comprised 1085 nodes in total. Nodes shown as circles along the edges were fixed nodes, including the entire third layer. The activity from the 10 sensors connecting to the mass at the center of the superficial layer (black node) were provided as input to the network. (**C**) Example sensory inputs used for network learning. One diagram illustrates the 10 sensor inputs for a Gaussian input, the other diagram illustrates the 10 sensor inputs for a skin input. (**D**) The 10 input locations used to apply the skin inputs are indicated on the first layer of the skin model, relative to the location of the central mass (black). Red dashed box below illustrates the skin sensor responses from the 10 springs attached to the central mass, and used as the training inputs, when one of the training inputs, Input #1 (Indent, Fig. S1), was applied to the different input locations (nodes colored in blue). The activity of all skin sensor signals were made to be positive-only by full-wave rectification. As an example, the skin input in panel C is the rectified skin input evoked from Position 5.

Another aspect that characterizes early developmental learning in infants and animals is random explorative behavior (22, 23, 24), which is quite different from the more repetitious training typically applied for machine learning networks. The former could be an advantageous strategy if the complexity of the underlying system that the brain needs to learn (i.e. the body and world dynamics in interaction) is so high that it is impossible to learn to represent it in full. Therefore, we trained the network on random sequences of non-repetitious activations of our previously published biological skin model, simulating a variety of dynamic touch conditions, where the sensors had extensive intrinsic dependencies (3). We found that the learned network could separate a diversity of spatiotemporal inputs that were unseen during the training (’naive inputs’) and also separate naive inputs with different kinematics. Ablation experiments after completed training showed that the network performance was still almost unaffected when parts of the sensor population or substantial parts of the network-internal connections were removed. Comparisons with the pre-training network across noise levels indicated that the training caused the network to better mirror the general dynamics of the sensor data from the skin model, where it can be noted that mirroring dynamics in effect is an emergent predictive coding strategy, since it forms a basis for predicting the next sensory input using preceding inputs. In principle, our results indicate that sensors embedded in a biomechanical system with intrinsic dependencies along with simple synapse-local learning rules can result in emergent learning effects that rapidly make the network operational by achieving initial, robust, ‘good enough’ solutions applicable to separate diverse unseen dynamic inputs to that biomechanical tissue.

## Results

### Experimental setup of the model

The biological motivations of the network component of the experimental setup is given in the Methods and the Introduction. Fig. 1A illustrates the network that comprised both excitatory and inhibitory neurons. Most of the experiments in this paper was done using a 10 x 8 network architecture, with 10 neurons in layer 1 (5 excitatory neurons and 5 inhibitory neurons) and 8 neurons in layer 2 (4 excitatory neurons and 4 inhibitory neurons), a total of 18 neurons, fully connected (’X’). Note that the terms ‘Layer 1’ and ‘Layer 2’ solely refers to neurons that received the sensory input and those that did not. These networks were trained to sensory inputs with and without intrinsic dependencies, using Hebbian-like plasticity across all synapses. Fig. 1B illustrates the mass-spring-damper model, which in a simplified manner emulates biologically quantified skin tissue dynamics during skin-object interactions (3). Its main purpose here is that it captures in principle the extensive dynamic co-dependencies between the mechanosensors that results from the complex mechanical couplings in the skin and in biological tissues in general. Fig. 1C illustrates one example skin input alongside one example Gaussian input, the latter being devoid of the sensor co-dependencies but matching the overall frequency and power distribution of the skin inputs (Fig. S2). The temporal profiles of the skin sensor responses illustrated that each sensor provided non-redundant information due to the distributed dynamic couplings.

In resemblance of the lack of behavioral structure that characterizes early learning in biological systems (22, 23, 24)), the network training consisted of a series of random, non-repetitive activations of the skin model, in what would correspond to less than one hour of random haptic interaction. We implemented randomness in the learning process by using a varied set of spatiotemporal force input patterns (Fig. S1) and by that every subsequent input was delivered to a random location on the skin (Fig. 1D). Because of the small size of the network, only the 10 sensors connected to the central mass (indicated in black in Fig. 1B, D) were providing input to the network, and their responses varied greatly depending on input location (Fig. 1D). Although it is beyond the scope of the present paper to provide an extensive account of the sensory information that arises in the skin on touch contact with external objects, here we provide a motivation for the stimulation patterns we used and how they should be interpreted in the wider setting of touch interactions. First, shear forces in the skin tissue is the critical factor that causes the skin mechanosensors to be activated (25, 26, 27, 3). The origin of shear forces is when the contact state dynamically changes, whereas static touch lacks information (28). Our stimulation patterns emulated a few different types of such dynamic state transitions, in terms of spatiotemporal surface forces (28, 26, 6) that subsequently dispersively propagate into the skin where they activate skin sensors in complex spatiotemporal patterns that outlasted the duration of the stimulus by a wide margin due to inertial effects in the skin tissue (3). The Gaussian nature (not to be confused with our Gaussian training inputs) of the structure of these specific skin stimuli contained a broad range of frequencies and each stimulus therefore in effect stimulated a wide range of dynamic properties in the skin model (3).

### Evolving network activity during learning

To test the impact of the intrinsic dependencies across the sensor population, we compared the learning obtained from skin inputs with the learning obtained in the same initial network instead trained on the pre-generated random patterns of Gaussian inputs (’Gaussian network’) (Fig. 2). During training, the inputs were provided in a continual fashion, which meant that the circulating activity within the recurrent network was never allowed to fully relax between consecutive stimuli, rendering each learning phase even more unique. Figure 2 illustrates the neuron activities comprehensively as well as in detailed snapshots, pre-(red arrows) and post-(green arrows) learning, for the ‘skin network’ as well as for the ‘Gaussian network’. On visual inspection, the neuron signals became more decorrelated and structured in the skin network compared to the Gaussian network (Fig. 2 C, D). The modeled neuronal responses to the spatiotemporal sensor input patterns in principle resembled those of synaptic inputs to intracellularly recorded cortical neurons *in vivo*, which also depend on various time-varying dynamics of recurrent connections that gradually unfold during the course of spatiotemporal sensory inputs (29, 30, 31, 32).

**Figure 2:**
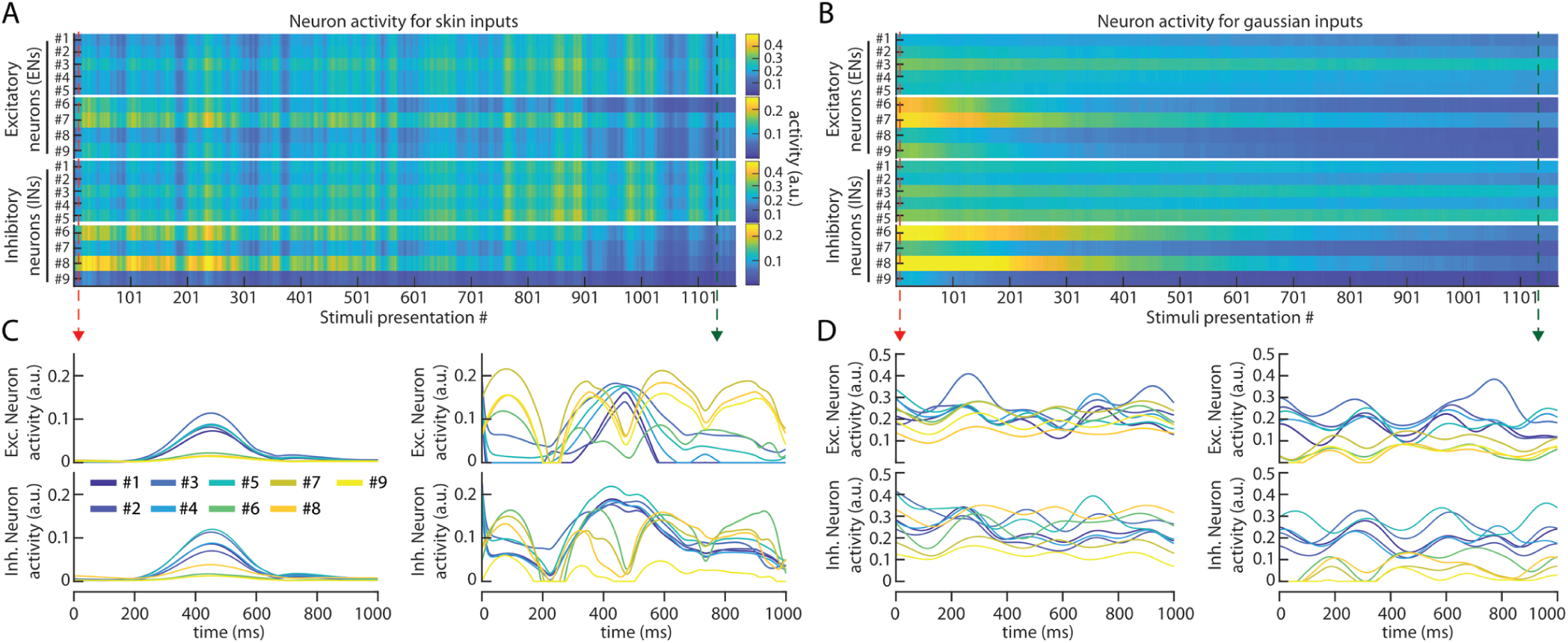
Evolving network activity during learning from skin and Gaussian inputs. (**A**) Individual neuron activities (moving average across 50 input presentations) during learning from an uninterrupted sequence of 1160 random skin inputs. This network had initial connectivity%=23 (see label definitions in text below). (**B**) Corresponding network learning from the Gaussian inputs. (**C**) Activities of the excitatory (top) and inhibitory (bottom) neurons (N=9 neurons per group, blue neurons were located in layer 1) to an example skin input at the beginning (left) and at the end (right) of the learning. (**D**) Similar display for a Gaussian input.

### Separation of naive inputs

A critical research question was if learning on the signals with the intrinsic dependencies (skin network) could lead to generalization in the network. We explored generalization by providing the trained network with inputs that it had not been exposed to during training (”naive inputs”, Fig. S1 F-J). To quantify the classification of naive inputs, we applied a linear classifier to the principal components (PCs) of the activity distribution patterns across the neuron population per each time step (N=3000) following the stimulus presentation.

Post-learning skin networks and Gaussian networks (Fig. 3A) were provided with naive skin inputs (Fig. 3B), which differed either with respect to their spatial force pattern (input type, Fig. S1) or they consisted of different temporal profiles of simple force impulses (i.e. with different kinematics) (Fig. S3). Qualitatively, for the five naive skin inputs, skin-trained networks exhibited more distinct neuronal population activity trajectories in PC space compared to Gaussian-trained networks (Fig. 3C; Fig. S4 illustrates the PC trajectories for the untrained, random network). Quantitatively, the linear classifier was applied to test the performance of five different initial network configurations (each with the same initial connectivity% of 23%, see section ‘Synaptic weight changes’) across a range of neuronal noise levels and found that the skin networks achieved a lower classification error rate for the five naive skin inputs compared to both Gaussian and pre-learning networks (Fig. 3D). The classification error rate was relatively consistent across all the different initial network settings (Fig. S5). When we tested the networks’ classification of force impulses that differed in their kinematics (Fig. S3;a variation of the input parameters that were not included in the training), the difference in both PC trajectories and classification performance between the two networks were even more pronounced (Fig. 3E,F).

**Figure 3:**
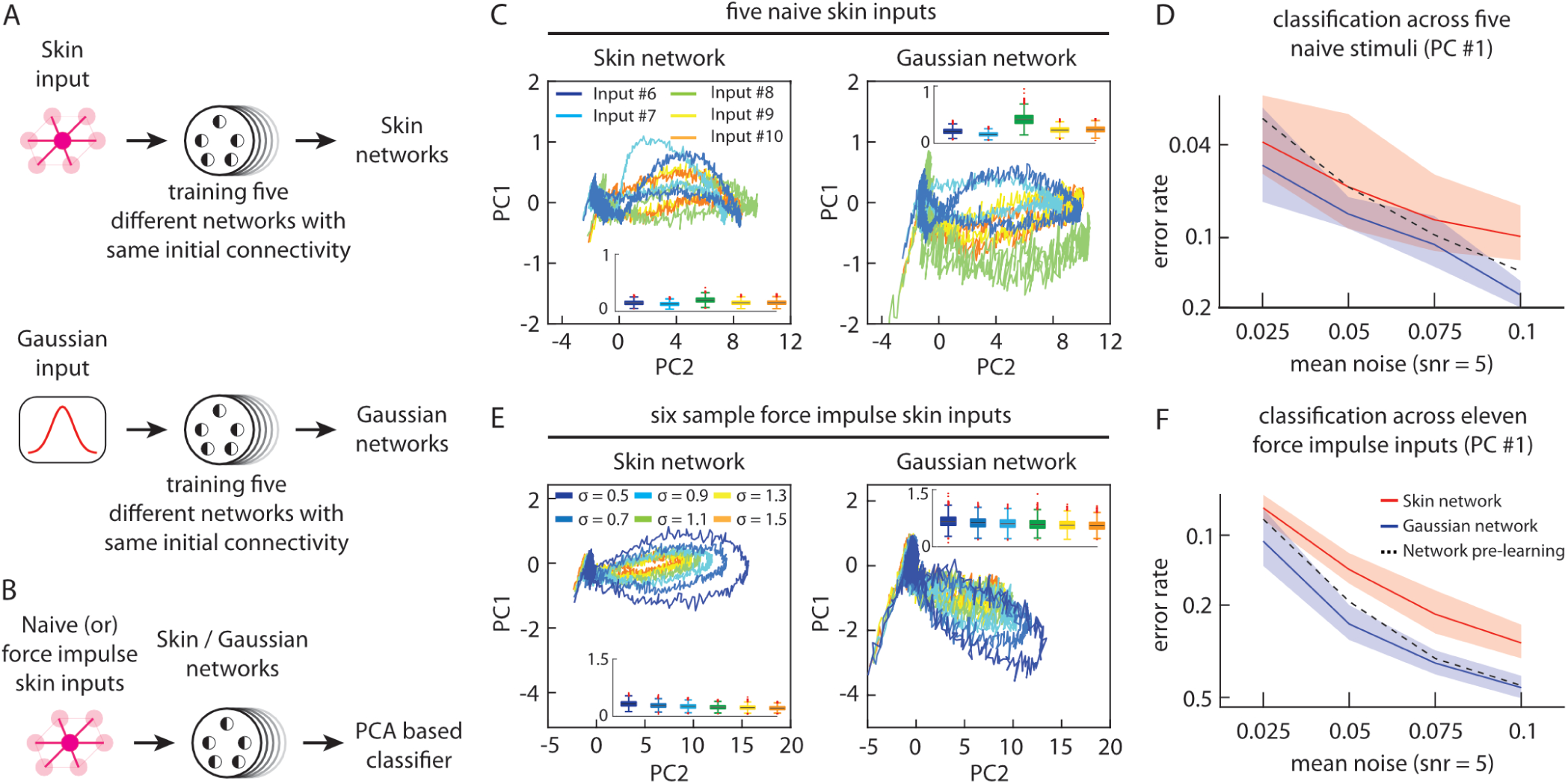
Generalization to naive skin inputs. (**A**) The network learned from skin inputs or from Gaussian inputs. (**B**) Protocols used for comparing the classification generalization. (**C**) The responses of the excitatory neuron population for the five naive skin inputs Fig. S1 F-J; color coded) projected on the first two principal components (PCs), comparing the skin and Gaussian networks (at a neuronal noise level of 0.1). *Inset*: the variances of the PC scores for the five different inputs. Same network as in Fig. 2. (**D**) Classification accuracies (mean and variance) across the five naive skin inputs for five different initial network configurations at the indicated noise levels. (**E**) Similar display as in **C**, but for eleven force impulse skin inputs (Fig. S3). (**F**) Classification accuracy across these eleven force impulse skin inputs.

Notably, across all tests, the skin network, unlike the Gaussian network, outperformed the pre-learning network, except at the lowest noise levels, where the networks approached deterministic behavior (Fig. 3E,F). The ability of untrained random networks to separate inputs during deterministic operation is consistent with the known behavior of RC networks. Such recurrent networks possess intrinsic dynamic modes that provide a capacity to separate dynamic inputs even without training of the weights of the intrinsic network synapses (33). However, neurons in the brain *in vivo* do not operate without noise (34, 35) and hence we believe our higher noise levels are more reflective of what biological neuronal networks can achieve in terms of input separation. The skin networks showed progressively better performance than pre-learning networks as noise levels increased (Fig. 3D,F), indicating that learning-induced adaptation of the intrinsic network structure based on inputs with sensory dependencies promoted network structures that focused on the dominant dynamics of those sensor signals. Hence, training on inputs with intrinsic sensor dependencies improved the networks’ generalized separation capacity, whereas training on inputs without such sensor dependencies (Gaussian networks) resulted in performance that was inferior to that of randomly connected pre-learning networks.

#### Network Scaling

As described in the Methods, the network architecture we used can be argued to resemble a primordial cortex, but the size of the network was of course substantially smaller than even in the earliest vertebrates. However, assuming that the learning processes has a decisive role in shaping the network connectivity, and thereby its emerging computational architecture, we hypothesized that the network size would not fundamentally alter the capacity of the trained network to separate naive inputs. In Fig. 4 we directly tested this idea by substantial increases in network size.

**Figure 4:**
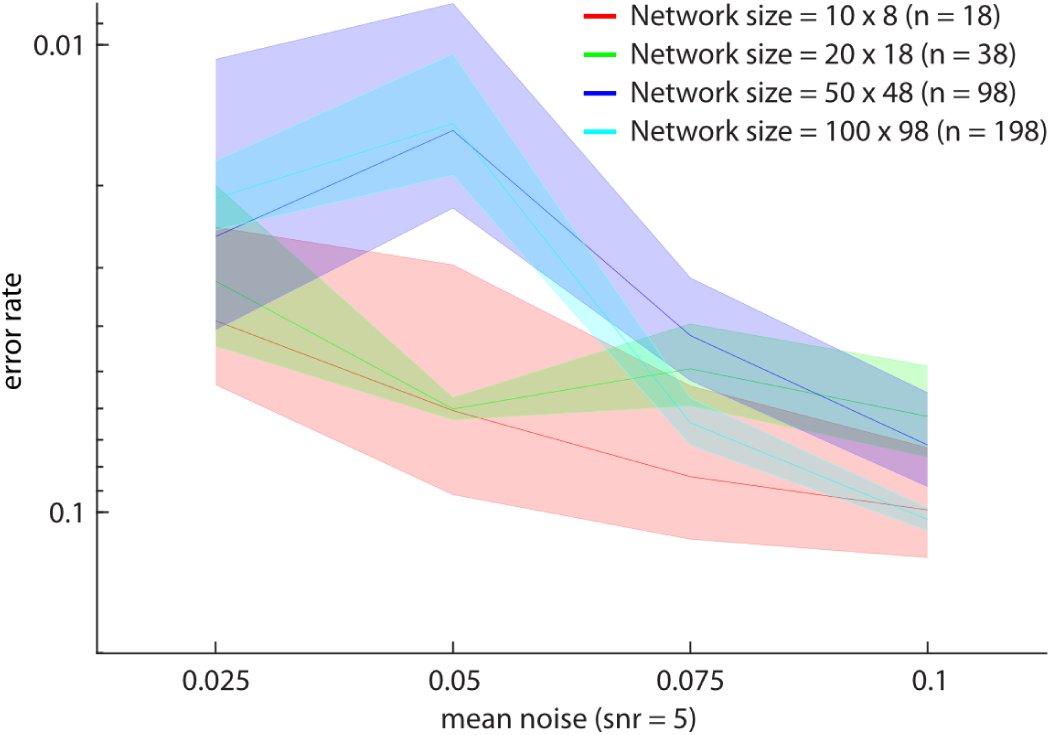
Classification accuracies (mean and variance) across the five naive skin inputs improved with the size of the network. Inset; ‘*n*’ indicates the number of neurons in each network size tested, *e.g.* the largest network had 100 neurons in layer 1 (50 excitatory and 50 inhibitory neurons) and 98 neurons in layer 2 (49 excitatory and 49 inhibitory), totaling 198 neurons, with full connectivity between all the neurons (symbolized by ‘X’).

When increasing the number of neurons in each layer, we kept the same connectivity rules for internal and external inputs. The classification performance improved slightly compared to Fig. 3D in larger networks, presumably due to better noise averaging from the increased number of neurons.

#### Robustness

Since biological systems can show a degree of robustness to tissue damage, sensor damage and even neural tissue damage, we also tested the impact of removing sensors and internal synapses post-learning (Fig. 5). Despite removing 20% of the sensors, the PC trajectories did not substantially alter (Fig. S6) and the classification performance for both naive and force impulse skin inputs (Fig. 5A) was unchanged or improved for the skin network (in comparison to Fig. 3D, F). In contrast, the Gaussian network showed reduced performance (Fig. 5A). Even more remarkable, the skin networks were highly robust to removal of network-internal synapses. Classification performance on naive skin inputs remained intact even when 75% of synapses were removed (Fig. 5B). This suggests that the training on the skin inputs induced systematic changes in the network structure, entraining the internal network activity dynamics to the dynamics of the underlying sensory dependencies, making the representations broadly distributed across the network and therefore highly robust. This would be conferring robustness to damages to the body, or to the brain (36), which would have major survival value in biology. As in Fig. 3D, F, the classification performance of the skin network progressively improved relative to the pre-learning network with higher noise levels (Fig. 5B), indicating that the learning of the skin network had resulted in a better approximation of the dynamic dependencies within the skin inputs. Learning on the Gaussian input again reduced the separation capacity relative to the pre-learning network (Fig. 5), an interesting effect that we will return to in the discussion.

**Figure 5:**
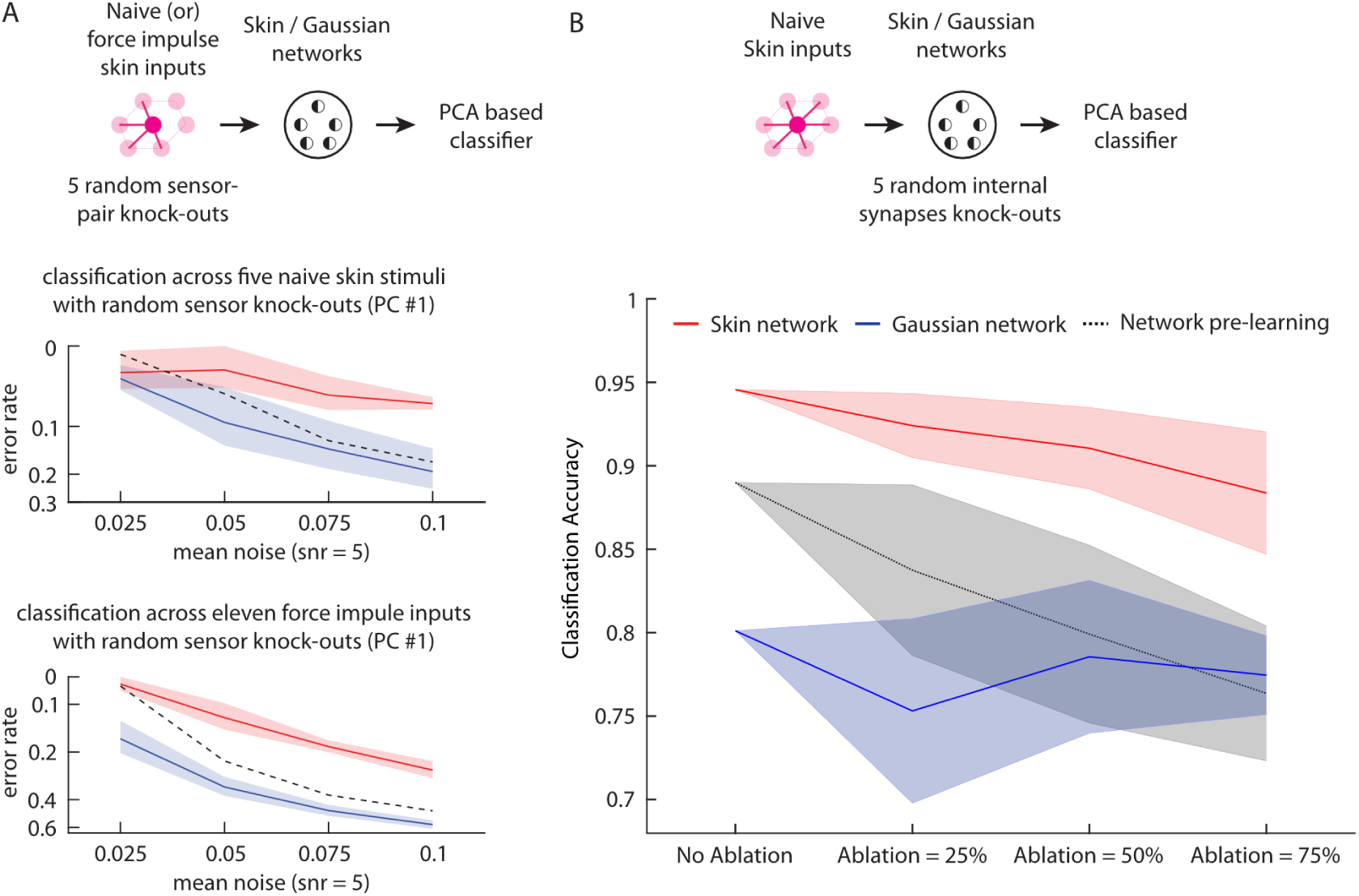
Ablation of random subsets of sensors or internal connections post-learning. (**A**) Protocol used for comparing the classification robustness with random sensory knock-outs. (**bottom**) Classification accuracies (mean and variance) across the five naive skin inputs and eleven force impulse skin inputs with random sensory knock-outs, at different noise levels. (**B**) Protocol used for comparing the classification robustness with network ablation. (**bottom**) Classification accuracies (mean and variance) across the five naive skin inputs, for three different levels of network internal synapse ablation. In the ablation study, 2, 4, and 6 randomly selected internal synapses per each neuron were removed from the network to simulate 25%, 50%, and 75% ablation levels, respectively.

### Synaptic weight changes

Because of their fully recurrent connectivity, even the smallest size of our network would exhibit complex system behavior. Complex systems are characterized by that the essence of their functional properties cannot be well described with mathematical tools. Therefore, to describe more precisely what changes the learning process resulted in within the network, we investigated the resulting synaptic weight changes. This was also interesting in relation to the existing neurocomputation literature, and the age-old credit assignment problem, i.e. can we ascribe any specific role to the changes in any single type of synapse, such as sensory versus internal synapses, internal synapses made on excitatory versus inhibitory neurons? Relative to the pre-learning network (Fig. 6A-C), training on skin inputs caused an increase in the weights of the sensory synapses (red dotted box, Fig. 6A), whereas the weights of the sensory synapses instead decreased in the networks trained on the Gaussian input (Fig. 6D; note the scale difference for the two networks). There were also extensive reorganization and excitatory/inhibitory re-balancing of the internal synapses (Fig. 6E), which gradually stabilized during the course of the training for both skin (Fig. S7, S9) and Gaussian networks (Fig. S8). This reorganization can explain how the skin network activity could decrease (Fig. 2A,B) when its synaptic weights increased.

**Figure 6:**
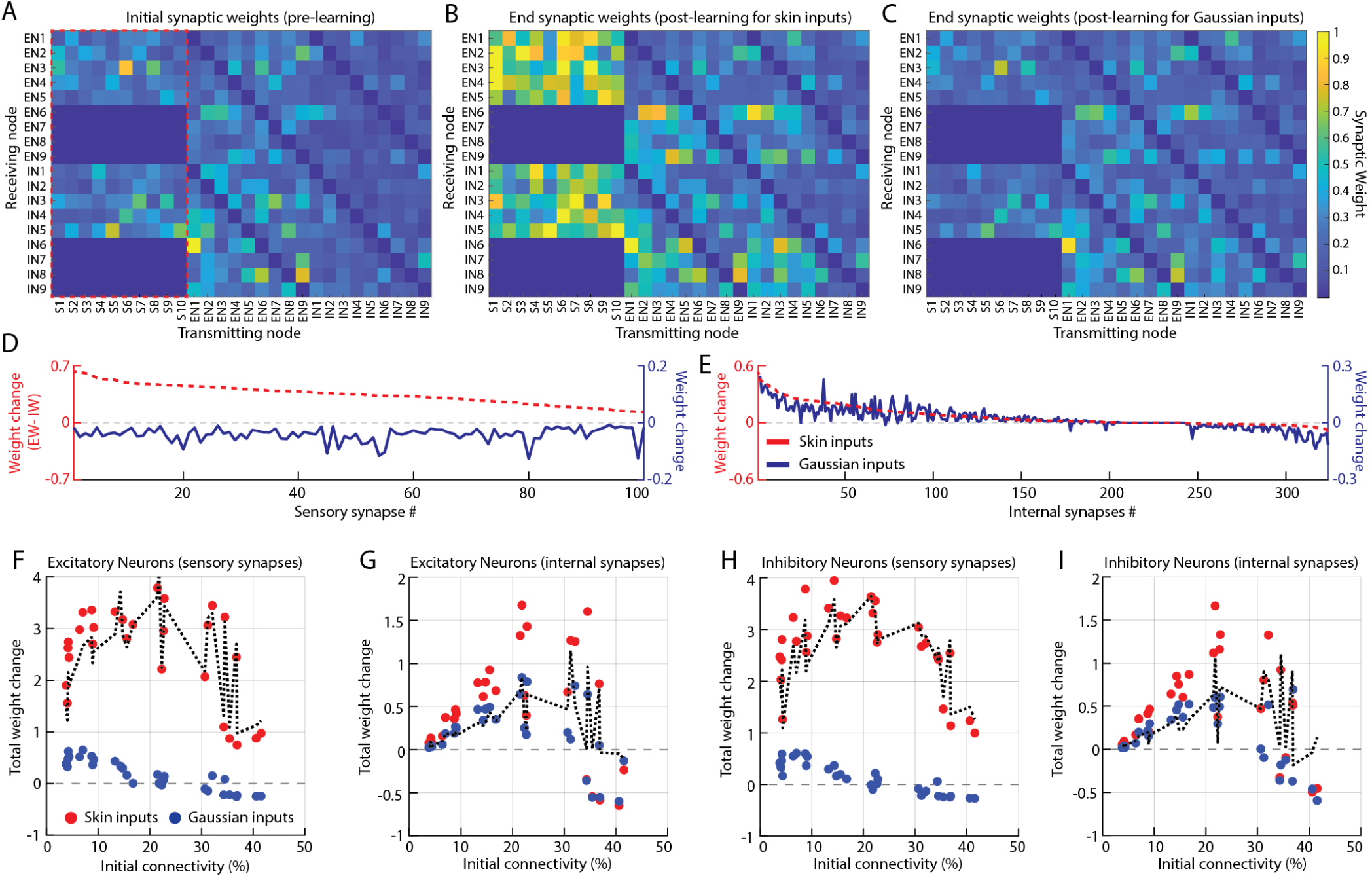
Synaptic weight changes following learning. (**A**) Initial synaptic weights for an example network (same network as in Fig. 1.) with the sensory synapses within the red dotted box. (**B**) The synaptic weights post-learning (end Weights) in the skin network. (**C**) The synaptic weights post-learning in the Gaussian network. (**D**) Sensory synapses sorted by their magnitude of weight change after learning from the skin input, for comparison of the skin network and the corresponding Gaussian network. (**E**) Similar display as in **D**, but for weight changes across non-sensory synapses (internal synapses). (**F**-**I**) Average net weight changes post-learning, sorted by type of synapse, across networks with varied initial weight configuration (Fig. S11, S12). The dashed lines indicate the difference in average net weight changes between the corresponding skin networks and the Gaussian networks.

We also explored the distribution of the synaptic weight changes more systematically across a variety of initial network configurations (Fig. 6F-I). To test the learning across a variety of initial network connectivities, we used 30 different network settings, each fully connected, identified by their ‘Initial connectivity%’. This label reflected the mean initial weights of the synapses but was also an index indicating the weight distribution (see Fig. S11, S12) and weight skewness (Fig. S13) when the training started. The initial network activity, which could be an additional factor impacting learning outcome, scaled roughly linearly with the ‘Initial connectivity%’s (Fig. S10), whereas the classification performance was overall comparable (Fig. S5). Fig. 6F-I illustrates that overall across all network configurations the sensory synapses changed the most, but the internal network synapses could change almost as much for the networks with intermediate Initial connectivity% (20-35%), in particular the feedback connections from layer 2 to layer 1 (Fig. S14). Further, the weight changes of the internal synapses on the excitatory and inhibitory neurons were similar across the different Initial connectivity%’s for the two network types (skin and Gaussian) (Fig. 6G,I). The skewness of the initial weights generally increased by the Initial connectivity% value (Fig. S13), but did not have any apparent relationship to these weight changes (Fig. S15). This analysis verified that across a relatively wide range of initial connectivity configurations, the learning resulted in comparable effects.

### The effect of response differentiation across the neurons

The activity of neighboring neocortical neurons exhibit a remarkable degree of decorrelation in the adult brain *in vivo*(37, 38, 39, 40, 31, 32). Because the emergence of such decorrelation is known to be beneficial for forming compact and efficient representations in the latent space of recurrent variational autoencoders (41), we also wanted to explore if this was an emergent effect in our networks. Therefore, we calculated the average cross-correlation, i.e. the correlation in the activity between two neurons at each time step across all time and across all pairs of neurons for each given network configuration (e.g. Initial Connectivity%). We observed that the cross-correlation measure decreased substantially with learning in the skin networks (Fig. 7A,B), indicating that individual neurons developed more independent activity patterns. However, for the networks with the highest connectivity% (Initial connectivity *>* 35%) such a drop did not occur, and these networks also had the lowest degree of restructuring of their network-internal synaptic weights (Fig. 6F-I). We hypothesized that the internal activity in these tight networks were too strong for their weights to become sufficiently impacted by the sensory synaptic activities. Therefore, we investigated if the low cross-correlation in these cases could be ‘rescued’ by a tweak to the learning rule such that the excitatory synapses were only allowed to ‘see’ a high pass filtered version of the postsynaptic signal used for learning. Implementing this ‘Excitatory synapse learning signal filter’ led to drops in the cross-correlations also within the networks with high initial connectivity%, with much lower impact on the networks with the lowest initial connectivity% (Fig. 7C, Fig. S16). Also the overall pattern of synaptic weight changes remained unchanged (compare Fig. S17 with Fig. 6F-I). Importantly, for the networks with high initial connectivity%, the learning signal filter did indeed also ‘rescued’ their input separation (Fig. 7D); it led to a reduction of the classification error rate from 0.12 (left) to 0.01 (right)).

**Figure 7:**
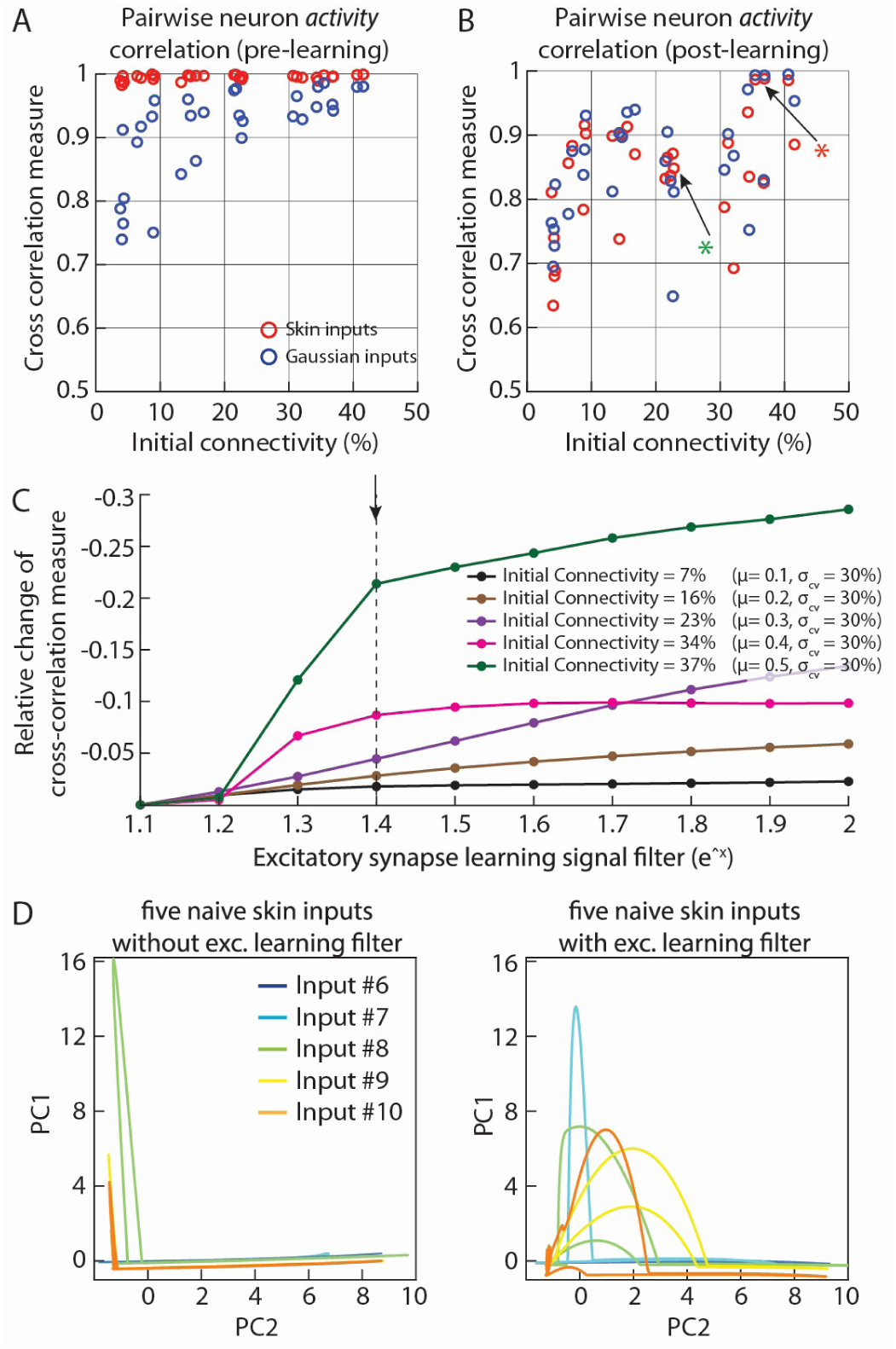
Diversification of the neurons’ activity patterns post-learning and ‘rescue’-effect of the excitatory learning signal filter. (**A**) Average pairwise cross-correlations across all the excitatory neurons pre-learning. (**B**) Same observation post-learning. Arrow with *green asterisk* indicate the network illustrated in Figs 2, 3 and the arrow with *red asterisk* indicate the network that failed to learn without learning signal filter. (**C**) Relative changes in the cross correlation measure in skin networks across different learning signal filter exponents. Dashed line indicates the exponent (1.4) used in the test in (D). (**D**) PC trajectories of the failing post-learning network without the learning signal filter (left) and the same network with the learning signal filter (right).

Finally, since for example inhibitory synaptic connections have been hypothesized to have different specific roles in the neurocomputation literature (see Discussion), we also investigated the impact of removing specific synapse types. Across all the tests we made, we found that removing inhibitory synapses, both laterally within the layers and the feed-forward connections, led to significantly higher cross-correlations after learning, whereas removing other connection types one-by-one did not have a consistent impact (Fig. S18). Hence these inhibitory connections could be more important for the network to acquire a high separation capacity.

## Discussion

Our paper was motivated by a lack of understanding of the network mechanisms that could make the early cortical circuitry operational. The assumption/research question was that learning from sensory dependencies could be at least part of the answer, a mechanism that can help lead to a bootstrapping of cortical circuitry functionality. We explored the emergent effects of Hebbian-like neuron-independent learning in a cortical-like recurrent network model when the sensory inputs were generated through an emulated biological skin tissue with complex mechanical couplings, which were activated in a non-repetitious, random manner, similar to early biological development (22, 23, 42, 43). Due to the sensory dependencies caused by the biomechanical couplings, this learning strategy rapidly converged towards generalizable network representations, applicable to separate previously unseen inputs with diverse spatial and/or kinematic characteristics. The network representations were also remarkably robust to post-learning loss of sensors and network connections, in addition to neuronal noise, all of which are beneficial effects in biological systems. Notably, the emergent functionality in our networks to learn to better mirror the dynamics of the skin model (3), has interesting parallels with findings in Reservoir computing networks which have been shown to have a potential to mirror and therefore predict specific dynamical system inputs field (33, 44). In the discussion below we also argue that the emergent property of the present autonomous intrinsic network learning to adapt the recursive network dynamics to the dynamics of the biological tissue is highly advantageous from an evolutionary perspective, where increased complexity in biomechanical properties of the body has been suggested to be a critical leading factor in the evolutionary race.

### What was the nature of the emergent network representations?

We showed that the network tended to learn the dominant dynamic behaviors of the skin model. This was supported by the conserved capacity of the network to separate the skin inputs when internal connections or sensors were removed after learning (Fig. 5A,B); and when increasing noise levels in the network led to that the skin network became increasingly better than the pre-learning networks (Figs 3,5). Training on input without the sensory dependencies (’Gaussian inputs’), instead degraded the network performance. Thereby, the specific dynamics caused by the mechanical couplings would tend to induce adaptations to the specific network dynamics, caused by the looping activity that the recurrent network connections will create (19). With autonomous learning, the aspects of the dynamic sensory dependencies that would be the most in concert with the network internal dynamics should be the most easily learnt. This is in line with recent observations from early cortical development (45, 21), where this principle was suggested to help the cortex to more rapidly become operational early in development. Since the dependencies across the sensors derive from the mechanical couplings in the tissue, which will impose a tissue-specific transformation on the sensory representation of any real world interaction, a similar set of dependencies would in principle be present across any stimulus/activation context. This could underpin a generalization capability i.e. separating or classifying touch-induced spatiotemporal force patterns that have not previously been experienced (Figs 3,5).

Notably, when the network begins to mirror the dynamics in the mechanical system, this is equivalent to the network beginning to predict or anticipate those dynamics (33). Hence, this learning strategy tends to emerge into predictive coding, which is widely held to be an important mode of operation in the cortex (46, 47). Predictive coding is a theory in neuroscience and machine learning suggesting that the brain constantly generates predictions about sensory input, comparing these expectations to actual data. Hence, any recurrent network that generates activity that to some extent is independent of the sensory inputs that it receives, but not in direct conflict with those inputs, will fit this definition. In the present paper, that generated activity was caused within the network itself, by the activity looping caused by the recurrent connections, when the network was excited by the sensory input.

In the predecessor to predictive coding (Dayan, 1995), it was noted that obtaining a good predictive function in that generated activity, one that mirrors or correlates with the input, is a difficult problem. Their solution was to guide network training by minimizing the entropy (or Helmholtz free energy) across a known set of data. In the present paper, the autonomous learning in the intrinsic synapses adapted the internal network dynamics to better match the dynamics of the biological tissue (as signalled by the dependencies between sensors), hence reducing the difficulty of the problem for the network to form predictions. This may make it more tractable for the brain to operate using predictive coding, in this case without requiring external guidance nor requiring any preformed hierarchy of layers in the network.

In contrast, training on the Gaussian inputs consistently degraded the performance compared to the pre-learning network. Since training on the Gaussian inputs did result in net positive changes in the intrinsic synaptic weights of the network, whereas the weights of the sensory inputs overall did not display a net change or even reduced (Fig. 6), it might be asked what was being learnt in this case? Here we believe that the network was excited by the unstructured sensory inputs such that the learning caused the network activity to become even more dominated by its own internal dynamics, which consequentially would reduce the matching between the dynamic modes of the network and the dynamic modes of the skin tissue model. At higher network ablation levels, the difference between Gaussian and pre-training networks was eventually lost (Fig. 5B). This suggests that the Gaussian input failed to induce deep structural changes in the network like the input with sensory dependencies appeared to do.

### Relationship to the literature

In our networks, the simultaneous excitatory and inhibitory synaptic plasticity would tend to lead to the inhibition canceling out the excitation. But as the sensory input is excitatory, inhibition will always be lagging ‘one step behind’, leaving some net excitation propagating through the network. This resulted in an emerging activity decorrelation between the neurons (Fig. 8, S18). In the networks with higher connectivity%, neuronal decorrelation did not evolve to the same extent, but was ‘rescued’ by the learning signal filter which also improved the input separation (Fig. 7). In biological neurons, the learning signal filter may for example be induced by that the dendritic spines, receiving excitatory synapses (48), differentiates the calcium signal available to the postsynaptic enzymatic machinery controlling the synaptic weights (8).

**Figure 8:**
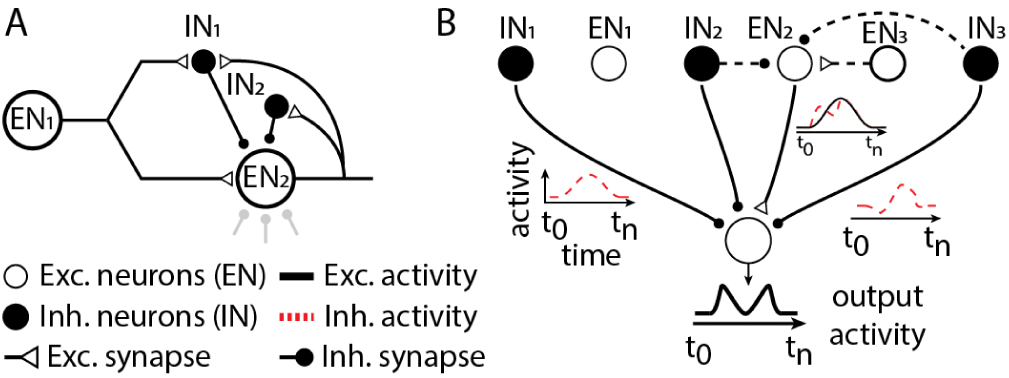
Diverse effects of synaptic inhibition in recurrent networks. (**A**) Scenario when feed-forward inhibition and feedback inhibition are mediated by the same inhibitory interneuron. Lower part, if multiple inhibitory interneurons provide input to *EN*_2_ (shaded synapses below *EN*_2_), the inhibitory control can sometimes have the properties of ‘detailed’ inhibition, and sometimes have the property of ‘blanket’ or ‘global’ inhibition (see text). (**B**) A feed-forward excitatory pathway via *EN*_2_ would lose its signal if the inhibitory signal was identical. A more diversified set of inhibitory signals (red traces) will transform and thus propagate parts of the excitatory signal (black traces).

Compared to the neurocomputation literature, We studied the plasticity-driven emerging functional roles for synaptic inhibition and synaptic excitation when there was no *a priori* constraint on their connectivity or what their learning should focus on (no ‘cost function’ or ‘error signal’). The neurocomputation literature instead often isolates excitatory from inhibitory synaptic plasticity during the learning process or considers inhibitory synapse learning as a homeostatic reaction to the learning in the excitatory synapses (12, 13, 14, 15, 16, 17, 18). It is common practice to regard the inhibition as having a specific role, such as feed-forward, feedback or blanket inhibition. More selective inhibition has also been modeled (49), in which case it could contribute to subsets of neurons contributing more independent computations (50, 10, 51, 52), in agreement with our decorrelation results above. But in a wider perspective, in recurrent networks it is hard to see how functionally ‘clean’ roles of inhibition could exist (Fig. 8A). A ground truth here could be that the division into excitatory and inhibitory neurons leads to that the inhibition can act to subtract aspects of the information (sensory or network-internal) that the network already ‘knows’ hence leaving ‘unknown’ information to be processed by other neurons in the network (Fig. 8B), both at the ‘global’ and at a more ‘detailed’ level, to use a proposed terminology (12).

The learning process described in this paper was a consequence of the contributions of all synapse types (Fig. S18), i.e. a consequence of the integrated network system. We do not claim that this is the best possible way to learn, only that these are the emergent effects we obtain under these biological-like conditions without any specific imposed learning goals. In relation to the machine learning literature, our networks share many overall features with various forms of RNNs. But more direct comparisons are difficult since the performance of recursive networks are notoriously difficult to quantify in terms of the types of meta-analyses that has been developed for feed-forward networks (53, 54), due to their partially chaotic nature (10), which is a consequence of their intrinsic complexity. We are not aware of any such meta-analysis for measuring the performance of fully recursive networks receiving complex population inputs with intrinsic dependencies. Instead, it is common practice in the application of recursive networks, including ESNs, liquid state machines (LiqSM) or RC networks, to simply randomize the internal synaptic weights of the network until a supervisor judges that some useful dynamic behavior emerges (33, 55) and learning is typically limited to the output connections outside this recursive network (11, 33, 55). Importantly, a given RC network can be controlled by external inputs to follow multiple alternative dynamic signals, as long as they are *a priori* known (44). In our network, the intrinsic connectivity was instead autonomously rearranging itself such that the network intrinsic dynamics found a better fit with a spectrum of dominant dynamics of the skin model, as signaled by the sensory dependencies. A recurrent network that autonomously adapts itself to dominant dynamic signals that are *a priori* unknown, may also be in a better position to separate previously unknown dynamic input signals generated via the same biological tissue dynamics..

### Relationship to models of brain operation

The proposed learning strategy aligns well with that most animals need to quickly learn to cooperate with the embedded dynamics in their body, because utilizing such *embodied intelligence* can lead to large gains in terms of control effort (4, 56, 57) and thereby shorten the time until the brain networks become operational. At the same time, a physical system such as the body with many components that interact with each other across various spatial and temporal scales is a definition of *complexity* in system theory. Indeed, the biological body can be considered to have evolved with respect to functional complexity, in order to increase the variety of actions the organism is capable of and thereby gaining advantages in the evolutionary race against other species (58). We believe that in order to utilize those advantages, it is crucial that the brain can adapt itself to whatever body it is situated in and to learn how to best mirror the dynamics that arise from those dependencies. Furthermore, because real-world interactions essentially are non-repetitious, it is the number of diverse, previously unencountered inputs that can be separated by the network (which we also referred to as network ‘solutions’ or representations) that would be the important performance metric. This is in contrast to performance metrics in machine learning which tend to focus on the precision by which any subset of given inputs can be identified. To evaluate our networks in the former regard we used PCA of the neuron population activity (Fig. 3), which is considered a well-suited analysis in the domain of complex systems (58). In this model of brain operation, the focus on a gradual emergence of ‘good enough’ (59) network solutions to the dynamics of the embodied intelligence, and separating diverse deviations caused by specific real-world interactions, could explain why the brain can become operational so rapidly but also why random explorative learning leads to a gradually improving control, as it does in infants (22). Notably, the network size, which for the cortex has increased through evolution, cannot be expected to alter the computational architecture *per se* but it would increase its capability to approximate and separate diverse sensory dependencies (Fig. 4).

In more general terms, it should be noted that the perceptual process is clearly very different from digital camera function, i.e. the cortex does not copy the information provided by the sensors but instead greatly transforms that input depending on its state (29). This can be understood by that the purpose of perception is to directly or indirectly drive behavioral choices, where behavior ultimately consists of spatiotemporal patterns of muscle activation (60). The neural activity patterns that drive that muscle activation are completely different from the sensor population input patterns, so one purpose of the cortex is to manage that transformation. We did not have any muscles in our model system, but the fact that the purpose of the cortex is to transform its sensory inputs remains, and that transformation and how it can gradually emerge is what our model is studying.

### Limitations

We aimed to keep the network as small and generic as possible, to facilitate interpretations of big picture functional effects, and to avoid assumptions regarding specific cortical connectivities. But this naturally prevented us from exploring the potential variations in the dynamic behavior in specific network configurations without full connectivity. Also, beyond the general functional principles that emerged, and that is in agreement with various biological data, at a detailed level we cannot validate our emergent effects using biological data, because there is no neurophysiological analysis available that can provide the required information on both single-neuron connectivity and the electrophysiological dynamic intracellular activity patterns across larger populations of cortical neurons.

Our ambition here was to provide a conceptual angle on fundamental emergent effects under a biological learning paradigm. Although our inputs were designed to be as complex and varied as possible within the constraints of the skin model, it could still be argued that we only tested these emerging effects for a set of relatively simple inputs, and that it remains to be seen whether the principle would stand in a more complex emulated body system with even more diverse interactions with a more complex simulated outside world. That would be a much larger endeavour. Our scope here was just to conceptually explore effects that would emerge, which is a new angle to approach the long-standing neuroscience issue of how cortical circuitry might become operational in its first early stages of physiological development. We hope that this can inspire tests under more complex conditions in the near future, in addition to inspiring new interpretational frameworks for neuroscience to better understand the mechanisms of how brain operation emerges from development.

## Materials and Methods

### Network Connectivity

We modeled the activity dynamics of two-layer, fully connected recursive neuronal networks comprising both excitatory neurons (ENs) and inhibitory neurons (INs) (Fig. 1A). In the standard networks, 10 neurons in layer 1 (5EN/5IN) & 8 neurons in layer 2 (4EN/4IN). Only the ENs and INs in layer 1 received external sensory inputs (see section *Sensory Inputs*) as excitatory synaptic connections, i.e., every EN and IN in layer 1 received 10 sensory inputs as excitatory synapses. The weight distribution of those sensory synapses are indicated under ‘Initial Synaptic Weights and Initial Connectivity%’ (below)). The networks were without autapses (self-excitatory/inhibitory connections) as they were previously observed to amplify the high-frequency parasitic oscillations within recurrent networks with these neuron specifications (19).

Our general network connectivity architecture, i.e. fully connected, was motivated from known features of subnetworks within the cortex. Notably, in its mature state the cortical network is not fully connected (61). But the rules deciding which connections are supported over time are not known and we chose to keep the network fully connected and instead let the emerging functionality within the network decide which connections should be reduced. And published data support that any neuron could in principle be connected to any other neuron. First, all cortical neuron types, excitatory as well as inhibitory are potential recipients of both thalamic inputs (62, 63, 64) as well as local and long range intracortical connections (63, 64). Hence, for the sensory input, all neurons of the cortex are potential recipients. But since our sensory input was derived from a small patch of skin, and since there is some level of topography of the sensory input even in the earliest vertebrate species (20), there would also be subsets of the cortical neuron population that would not receive input from that particular skin patch. Hence, parts of our neuron population did not receive the sensory input (referred to as ‘Layer 2’ neurons). Our fully connected network was motivated by that recurrent excitatory loops are a common feature in neocortical circuits (65, 66, 67, 68, 69, 70, 71), that inhibitory interneurons provide lateral inhibition (8, 72, 73, 74) as well as feed-forward inhibition (63, 75), but also make synapses on other inhibitory neurons, thereby also potentially forming recurrent disinhibitory loops (76, 77, 78, 79). Hence, both excitatory and inhibitory synapses can form all-to-all local cortical networks (71) in addition to providing long range intracortical connections (80, 81, 82).

### Neuron Model

Our previously published non-spiking linear summation neuron model (LSM) was used to compute the activity of both excitatory and inhibitory neurons. LSM is a compact model that emulates the impact of membrane time constants and conductance-based synaptic integration in biological neurons. It is hence capturing the central characteristics of an H-H conductance-based neuron model (19) where the time-continuous output activity is intended to reflect the mean firing rate of a traditional spiking neuron model. In a biological neuron, due to that its electrical signaling is being driven by opening and closing of conductances (ion channels), the impact that the activation of a given synapse results in does not only depend on its synaptic weight, but also on how many other conductances are open at the same time. The LSM represents two types of such conductances. First those of other synapses active at the same time. Secondly, the static leak conductance, which represents the constitutive leak ion channels that establish and maintain the resting membrane potential and that in the biological neuron therefore needs to be always open. A natural consequence of this conductance-based signaling, just like in the biological neuron, is that the higher the activity of all other synapses (or conductances) on the neuron, the lower the impact of a given synapse with a given weight (i.e., a given conductance). This can help protecting neurons from overexcitation, for example, and thus can contribute to stability in the learning network (19). In the LSM, the membrane potential (*A*, Eq. 1) of a neuron is given by the summation of weighted (*w*) input synaptic activity (*a*), that is normalized by emulated membrane conductances (static leak and dynamic leak). Here, *w_i_* and *a_i_* represent the weight and activity of synapse *i* (Eq.1). The static leak (*k*) emulates the leak channels contributing to the resting membrane potential and is scaled by the total number of synaptic inputs (*n*) to the given neuron (*k* = 0.25 per synapse). The dynamic leak (*τ*) reflects the rate of leak towards the resting potential, normally limited by the membrane capacitance of the neuron, which was a constant (*τ* = 1*/*100). The scaling effects of these membrane constants on the neuron activity under various synaptic input conditions were previously reported in (19). The neuron output activity was thresholded at a membrane potential corresponding to zero net synaptic input activity in the neuron (Eq. 2). The neuron activity of LSM is given by Eq. 1 & 2.

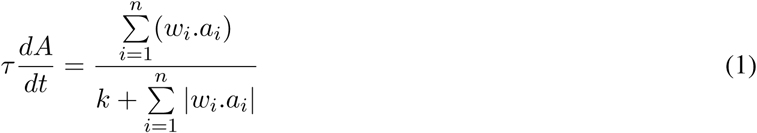

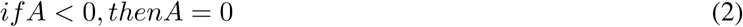

### Sensory Inputs

#### Skin Model

We used a mass-spring-damper model of a slab of skin (Fig. 1B) to generate the sensory input signals (*skin inputs*). This skin model was previously developed to replicate the dispersive mechanical waves that are elicited in biological skin during tactile interaction (3). The spreading mechanical waves, also studied in other experiments (83, 84, 85) create differential skin tissue movements, hence generating shear forces (26, 3), that are known to activate the skin mechanosensors. The masses (circles, Fig. 1B) in this skin model were arranged in three layers, superficially resembling the structural properties of human skin (epidermis, dermis and subcutaneous tissues). The masses were homogeneously distributed in a mesh-grid-like structure along each layer, and in pyramidal structures across layers (color coded as per layer, Fig. 1B). The neighbouring masses were interconnected via a network of springs and dampers (lines, Fig. 1B). In this study the skin model has a mesh size of 20×20 masses in layer 1 (400 masses), 19× 19 masses in layer 2 (361 masses) and 18 × 18 masses in layer 3 (324 masses); interconnected with a 5770 springs and dampers. The model uses the inter-node distance, which is a representation of strain, corresponding to the spring load, as the best available proxy for representing the signal that would drive the activation of mechanoreceptors. The masses on the edges of the skin model (empty circles, Fig. 1B) were fixed in all three axes. The equation of motion for a spring-mass-damper system with multiple degree-of-freedom is given by Eq. 3 (86).

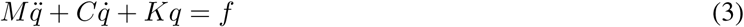

Where *M*, *C* and *K* are matrices denoting mass, stiffness and damping of the whole system and *f* is the input force vector. *q*, *q̇* and *q̈* are denoting position, velocity and acceleration of the system respectively. In order to numerically integrate and simulate the equation of motion (Eq. 3), it must first be reformulated as a state-space model (Eq. 4). A state-space model provides a linear representation of a dynamic system, making it suitable for numerical computation (86, 3).

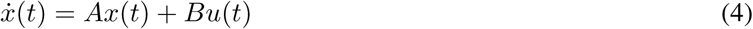

Where *x* is the state vector, *u* is the applied input force, *A* is the state matrix, and *B* is the input matrix. Coupling equation 3 and 4, we obtain

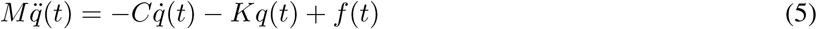

Which can be written as,

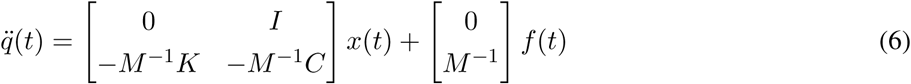

To compute the numerical solution of the above-described state equation we used an *ode45* solver (fifth order Runge-Kutta method) in MATLAB, with a time step of 0.01 ms. Each mass within the skin model moved freely in the three-dimensional space, the matrices and vectors in the above defined state equation were adapted accordingly. The values of *M, K*&*C* are 1.0 mg, 2.5 N m^−1^ & 5x10^−4^N s m^−1^, as studied in (3).

#### Skin Inputs

To activate the skin sensors, we designed different spatial skin force patterns that were activated by defining the input force vector (*f*) of equation 6 as a Gaussian function with a sigma of 1*ms* (Fig 1B, insert). The force vector was three dimensional (*f* = [*f_x_f_y_f_z_*]), describing the input force across all three axis of the skin. The obtained force across each spring was used as the as the output signals of the skin model, used to drive the activity of the neuronal network (*Skin Input*, Fig. 1C). The forces were given by Hooke’s law (*F_s_* = *kx*), where *k* is the stiffness constant of the spring and *x* is the rate of change in the position from equilibrium. Out of the 5770 springs available in this skin model, we used only the ten springs connected to the mass in the center of layer 1 (black mass, Fig. 1B). The responses of these ten springs were used for training and testing the networks. The activity of all skin sensor signals were made to be positive-only by full-wave rectification of the spring tension signal.

#### Gaussian Input

As a comparison for the networks that learned from skin model inputs, which have systematic intrinsic dependencies or sensor population regularities across all inputs, we also let some networks learn from ‘Gaussian inputs’ that lacked such intrinsic dependencies. For each of the 10 Gaussian ‘sensors’, we used pseudo-random spike trains with a spiking frequency of 1-3 Hz frequency drawn from a normal distribution and convoluted using a Gaussian function to obtain a time-continuous sensory input (*Gaussian Input*, Fig. 1C). An inbuilt MATLAB function “*randi*” was used to generate the spike time distributions and “*gaussmf*” was used for the convolution of the spikes into time-continuous responses. The distribution and convolution parameters for generating the Gaussian activity were chosen to match the power and frequency components of the skin inputs (Fig. S2).

### Network Training

For learning we provided the network (Fig. 1A) with a series of 1160 stimulus presentations, either sensory inputs from the skin model or from the Gaussian model (Fig. 1C). A network that learnt from the skin inputs is referred as “*skin network*” and a network that learnt from the Gaussian inputs is referred as “*Gaussian network*”. For the skin networks, we used 50 unique skin activation conditions, consisting of five different input types (*indent, push, pull, slide, point push,* Fig. S1) at ten different locations on the surface of the skin model. The 50 different inputs were provided in a predefined (pseudo-)random order. For the Gaussian networks, we generated non-repetitive random activity patterns across the 10 sensors. For both skin and Gaussian training inputs, individual stimulations were applied in a pseudo-random order, at an interval of 3000 ms, for a total of 1160 repetitions, and a total of less than 1 hour of training. Notably, predefined pseudo-random orders were applied to both the skin training and the Gaussian training so that the effects of changing other parameters of the network (such as seed connectivity) would be comparable.

### Initial Synaptic Weights and Initial Connectivity%

All the initial synaptic connections (”*initial weights*”) in the network were given a weight between 0.001 and 1 (*initial weights*). No weight could ever exceed 1. As all our networks were fully connected no synapse had 0 weight. The initial weights of the whole population of synapses in the network were generated as log-normal distributions with different mean weights (*µ*) (values between 0.05 to 0.5) and coefficient of variation (*σ_cv_*) (values between 10% - 50%). The variance of the distribution was quantified as (*σ*) = (*σ_cv_/*100) ∗ *µ* (Fig. S11). ‘Initial connectivity%’ indicated the mean weight in the network but was also a label we used to indicate the different initial weight distribution parameters (*µ* and *σ_cv_*) (Fig. S12). Within each given parameter combination (initial connectivity%) weights were also randomized to obtain five different versions of each initial connectivity% network, which were used to obtain the variances indicated in Figs 4-6.

### Synaptic Learning

We used the same Hebbian-inspired ‘*activity dependent synaptic plasticity rule*’ (8) for learning synaptic weights in both excitatory and inhibitory synapses. The learning rule was formulated to reward the correlation between the individual synaptic activity (*a_i_*) and the neuron activity (*A*). A positive correlation between the neuron & the synaptic activity leads to potentiation (increase) in synaptic weight and decorrelation between these activities leads to depression (decrease) in synaptic weight. For each stimulus presentation (*t*_0_ − *t_n_*), the weight change (Δ*w*) of a synapse (*i*) was given by equation 7.

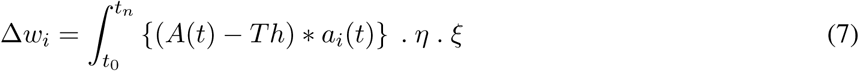

The value of the integral (*t_n_* = 1000*ms*) determined the magnitude of the individual synaptic weight changes.

To obtain long-term network stability, we also implemented a regulation of the synaptic plasticity that is related to the BCM rule, using a *learning polarity threshold (Th)*. The learning polarity threshold implements an effect that is related to the BCM rule and defines above what activity level the neuron becomes more likely to depress its afferent synapses, and vice versa (87, 8, 88). The *Th* was given by equation 8. In this case, we did implement a minor difference between the excitatory and the inhibitory synapses, which led to that the learning polarity threshold was lower for the inhibitory synapses at each given activity level of the postsynaptic neurons (Fig. S19C). This means that the probability of potentiation in the inhibitory synapses was higher for each given neuron activity level, although to a decreasing degree as the neuron activity increased. In particular at low-levels of neuron activity, such as in early stages of learning, this non-linearity enhanced the propagation of excitatory activity through the network.

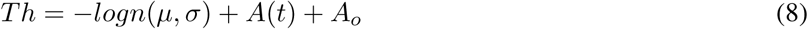

The strength of potentiation/depression of an individual synaptic weight was also regulated by a learning rate (*η*) and a compensation factor (*ξ*). The learning rate was defined as an exponential decay function (*e*^−3.5^) of the current neuron activity, limiting the rate at which synaptic weight changes occurred (Fig. S19A). The compensation factor constant is given by a sigmoid function (midpoint of 0.5 and steepness of 0.05, Fig. S19B) of current synaptic weight *w_i_*. The compensation factor allowed the weight change of a given synapses to become higher if its current weight was low and vice versa. This factor is motivated by that the insertion of a synaptic receptor in the synaptic membrane can be regarded as a chemical reaction (the insertion and removal of synaptic receptors is handled by partly separate chemical reactions (89)), where the addition of a new channel will become less likely the higher the number of ion channels already inserted and vice versa (19). The compensation factor function was flipped for depression and potentiation of a synapse (Fig. S19B). The weight change (Δ*w_i_*) and learning polarity threshold (*Th*) were computed as moving averages across 100 stimulus presentations in order to dampen their rates of adaptation.

#### Excitatory Synapse Learning Signal Filter

In general, the excitatory and inhibitory synapses learned using the same Hebbian-like learning rule, based on their activity correlation with the postsynaptic neuron. But for the densest network, this proved to not work as well as for the less densely connected networks, and we therefore implemented a tweak where the excitatory synapse learning was instead based on correlation with a high-pass filtered version of the postsynaptic neuron activity. The purpose was to make the excitatory synapse learning more selective, i.e. it would only correlate with positive changes in the postsynaptic activity. This could for example equal that the the postsynaptic activity in the dendritic spine, or the synaptic subspace underneath the excitatory synapse, would only be exposed to or only react to positive *changes* in the calcium activity of the main compartment of the neuron. Biological studies indicate that the near-synaptic calcium concentration can have different time constants than the calcium concentration in the main compartment of the neuron (90, 91, 92) and specific dynamics in the calcium concentration could impact synaptic plasticity (87). In order to achieve a high-pass filtered version of the postsynaptic activity (*a_i_*), we multiplied the synaptic activity by an exponential function (*e^x^*), as shown in equation 9. We tested the effect of synaptic weight learning across different orders of that exponent (Fig. 5C). This filter was applied only on excitatory synaptic activity (*a_i_*) during the learning phase, in the analysis of Figures 5C,D, S16, S17.

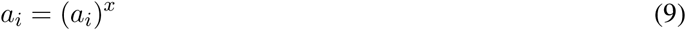

### Network Performance (Post-learning)

To evaluate the learning performance, we compared the generalization and the robustness capabilities of the skin networks and the Gaussian networks (Fig. 3). In order to test the generalization, we quantified the classification accuracy across five naive skin stimuli (Fig. S1F - J) or 11 force impulse skin stimuli (Fig. S3), neither of which were used during the learning process (Fig. 3D, F). These inputs were in this testing phase applied to the central mass only (Fig. 1). In order to test the robustness we quantified the classification accuracy across the naive and force impulse input, by randomly removing two sensor input synapses, corresponding to 20% of the total sensor input to the network.

The classification performance was quantified for four noise levels, where random noise of a given signal-to-noise ratio (*snr*) was added to the activity of each neuron. For each network configuration tested, we reported the average classification performance across the test inputs with 5 different repetitions of each noise level (Fig. 3,4,5).

### Statistical Analysis

#### Total Weight change

Total weight change (Δ*W*) within a neuron is defined as the summed difference between all the initial weights (W*_i_^I^*, pre-learning) and their respective end-weights (W*_i_^E^*, post-learning) as given by equation 10. Where, *m* is total number of neurons (*j*) and *n* is total number of synaptic weights (*i*) for a given neuron. The average synaptic weight changes across all the neurons was reported in Fig. 4F-I, S14, S15, S17.

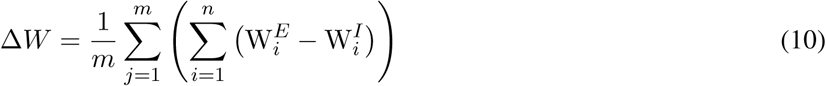

#### Cross-correlation

The correlation index measure was used to compute the similarity in the neuronal activities during test inputs. The correlation between two neurons was computed with an inbuilt MATLAB function “*xcorr*” (with zero lag), which produces values from 0 (uncorrelated) to 1 (identical). This pairwise cross correlation was calculated across all neuron pairs and the average value was reported (Fig. 5A-B, S18, S16).

#### Frequency analysis

A continuous wavelet transform (using an inbuilt MATLAB function “*cwt*”) was performed in order to define the frequency composition of the input sensory responses over time. The wavelet transform was used to extract the power of each frequency band as a function of time for the time-continuous responses of the sensors. Here, we reported (Fig. S2), for each frequency band, the maximum power of each sensor across ≈ 1160 stimulus presentations.

#### Principal component analysis (PCA)

Principal component analysis was performed on the activity of the excitatory neurons using an inbuilt MATLAB function *pca*. The pca function projects the neuron activity onto principal component space (new set of variables, Principal Components (PCs)) that explain the high dimensional input data on fewer output dimensions while encompassing the maximum preservation of information (variance) in the input data. The product between the neuron activity and PC coefficients results in “*PC scores*” that represent the sensory signals in PC space (Fig. 3C, E).

#### Classification

To quantify the classification accuracy across different skin inputs (Fig. 3D, F) we used a linear tree classifier. The excitatory neuron population activity for each given skin input at each defined level of network noise was projected onto the first PC and the PC scores (see section above) were used as input observations to the classifier. Each skin input was repeated for 50 times, where the neuronal noise resulted in different responses for each repetition. The data to the classifier was binned into 5 classes while testing the 5 naive skin inputs (50 repetitions x 5 classes = 250 observations), and 11 classes while testing the 11 force impulse inputs (50 repetitions x 11 classes = 550 observations). As each response consisted of 3000 time steps, the classifier had 3000 predictors for each observation. The classifier was trained and tested using a 5-fold cross-validation, which was repeated for 100 iterations to ensure robustness. This analysis was performed using a Matlab toolbox “*Classification Learner*”.

## Data Availability

The full set of simulation data used to produce all of the results in this paper is available from the authors upon reviewers request and at publication.

## Acknowledgements

This work was funded by EU H2020 FET Open project no. 829186 ‘ph-coding’.

This work was supported by the EU Grant FET 829186 ph-coding (Predictive Haptic COding Devices In Next Generation interfaces), the Swedish Research Council (Project Grant No. K2014-63X-14780-12-3).)

## Author Information

### Contributions

U.B.R. and H.J., designed and performed the research and wrote the paper. U.B:R: conducted the work and the analysis.

## Supplementary Material

**Supplementary Figure 1:**
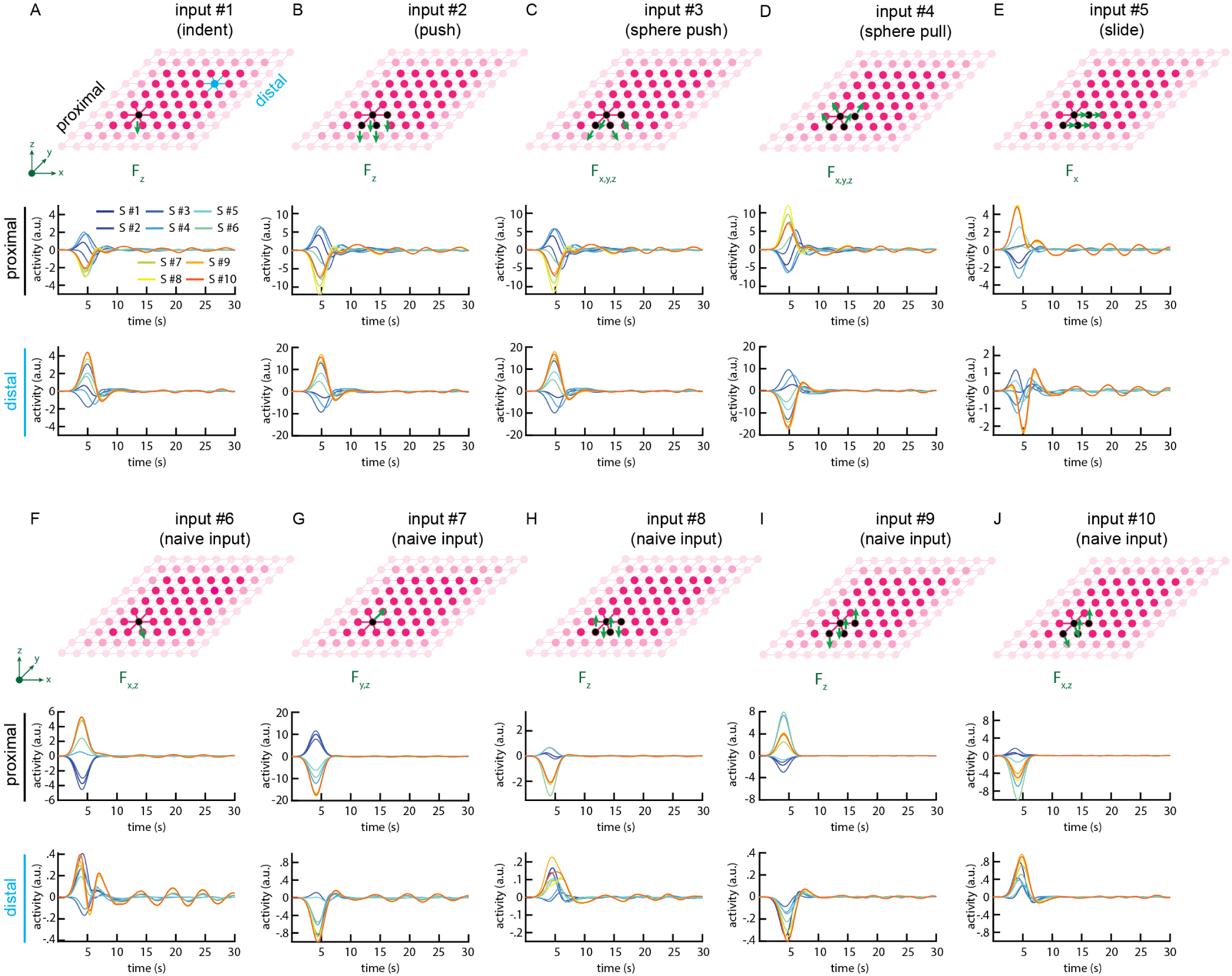
Skin sensor responses (spring forces) from the 10 springs attached to the proximal (node colored in black) (used as S#1 - S#10 for the network training) as well as 10 springs attached to a distal node (node colored in blue). The responses of the two population of sensors were shown separately for each input. Note that each input had the same duration but differed with respect to the spatial force distribution along the x-/y-/z-axes ([*f_x_f_y_f_z_*], as indicated by the green arrows). (**A-E**) The five stimuli used for network learning. (**F-J**) The naive stimuli used for testing generalization capabilities in Fig. 3. Note that the actual skin model was composed of more nodes than illustrated in the display.

**Supplementary Figure 2:**
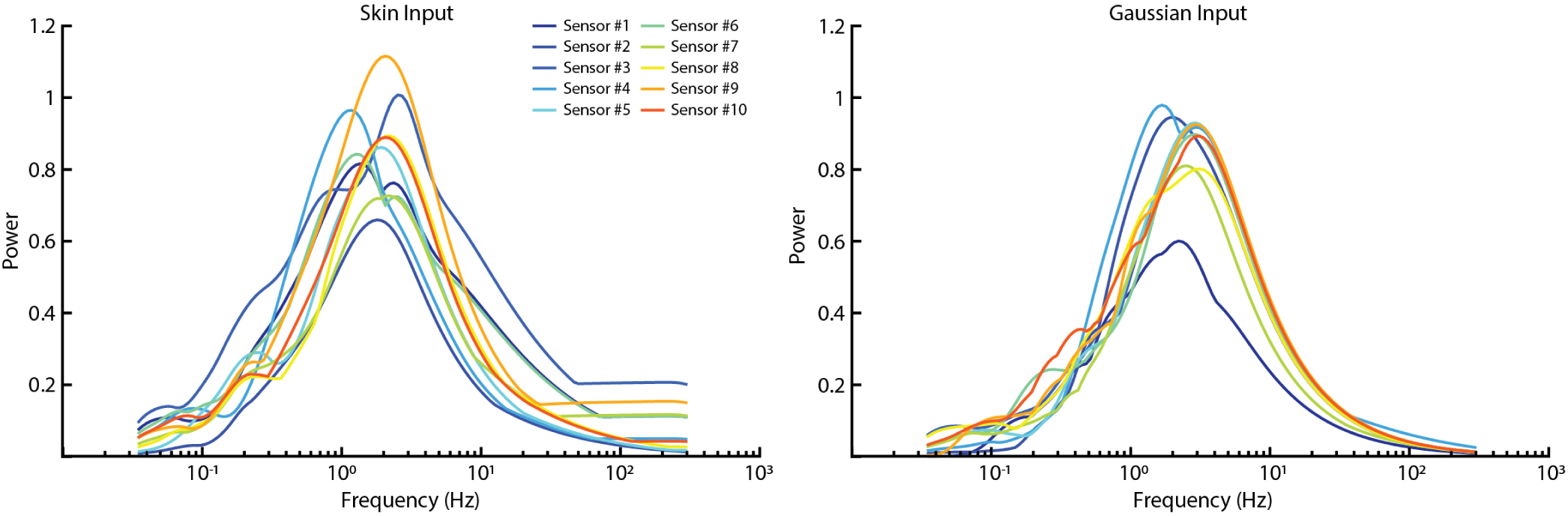
Frequency power distribution of all ten skin and Gaussian sensory inputs, averaged across 3500 stimulus presentations.

**Supplementary Figure 3:**
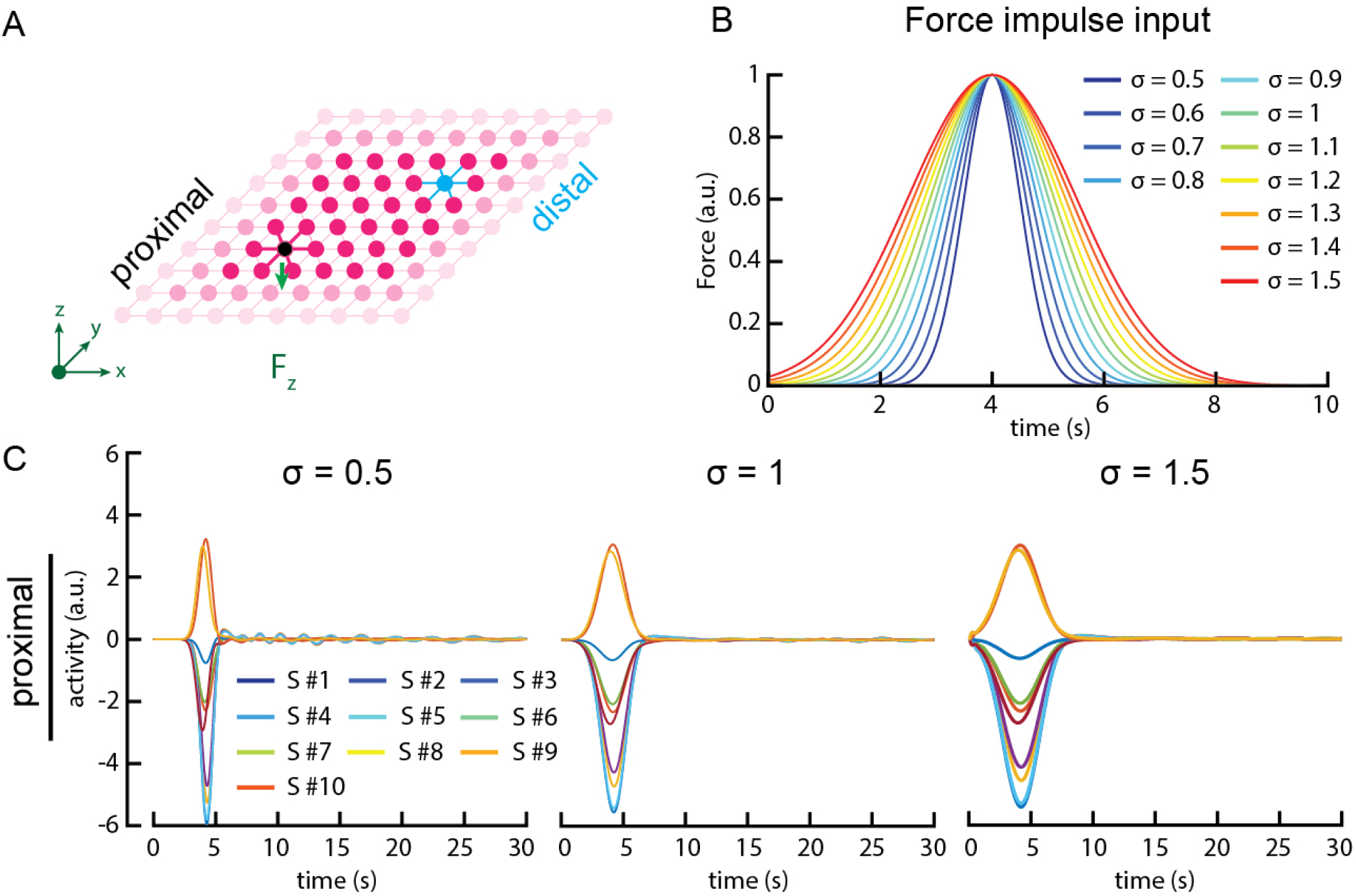
The force impulse input consisted of applying a time-evolving force to one mass only. The mass in black was impacted by a force in the z direction (A). (B) The force was applied with a Gaussian function, and force impulses differed solely by the sigma of their temporal Gaussian function. This contrasted with the skin inputs for which the temporal Gaussian function was always the same (*σ*=1). (C) Example responses of the proximal sensors, i.e. the 10 sensors connected to the proximal mass.

**Supplementary Figure 4:**
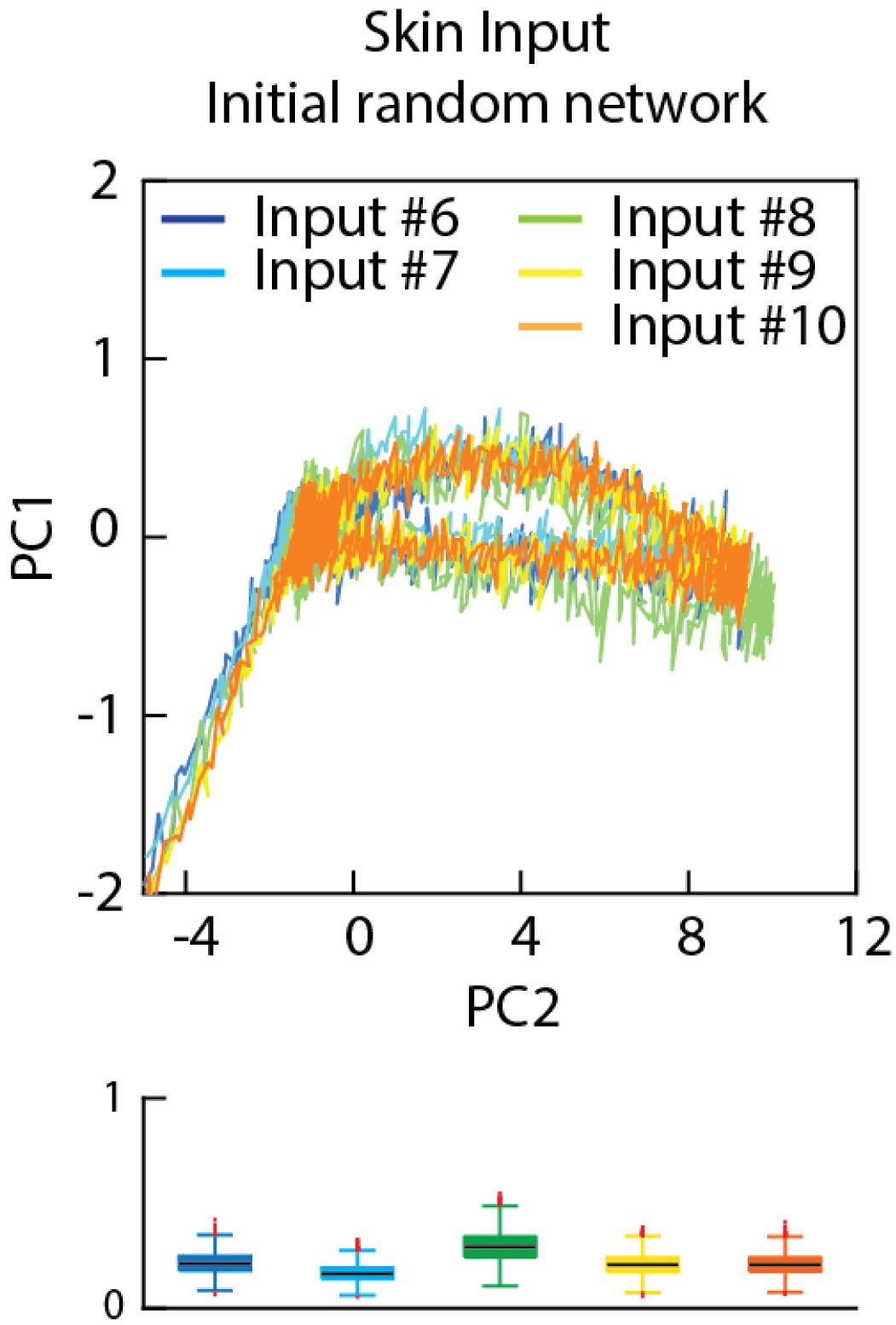
Projection of the neuron population activity from the pre-learning random network onto principal component space. The output of the neuron population for five different naive skin inputs (Fig. 1F-J) projected on the first two principal components (PCs) (at a neuronal noise level of 0.1, same as in Fig. 3C,E). Inset: the variance of the PCs across the 50 different initial noise randomizations.

**Supplementary Figure 5:**
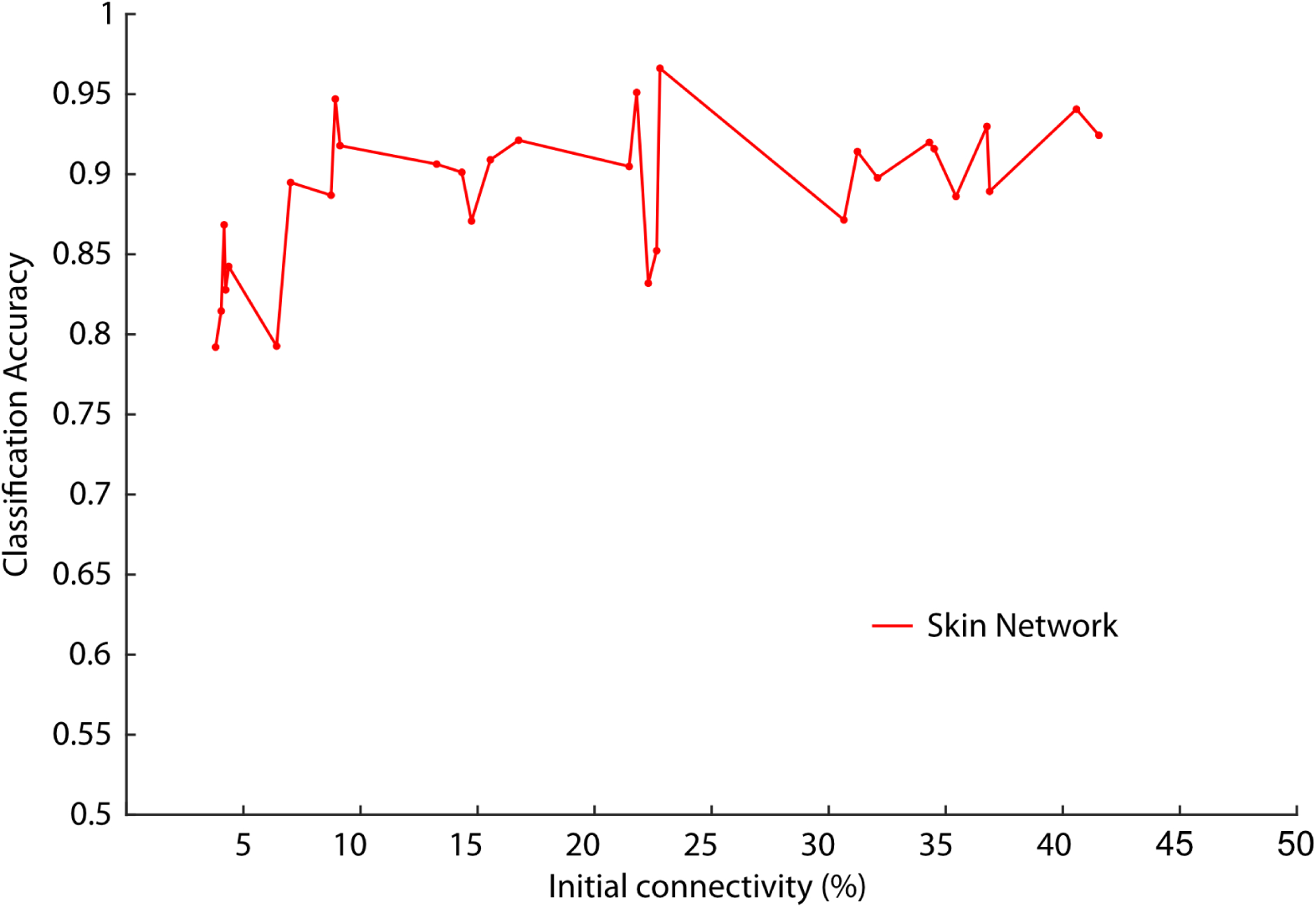
Classification accuracy across the five naive skin inputs (similar to Fig. 3) for Skin networks with varied initial connectivity (at a neuronal noise level of 0.1).

**Supplementary Figure 6:**
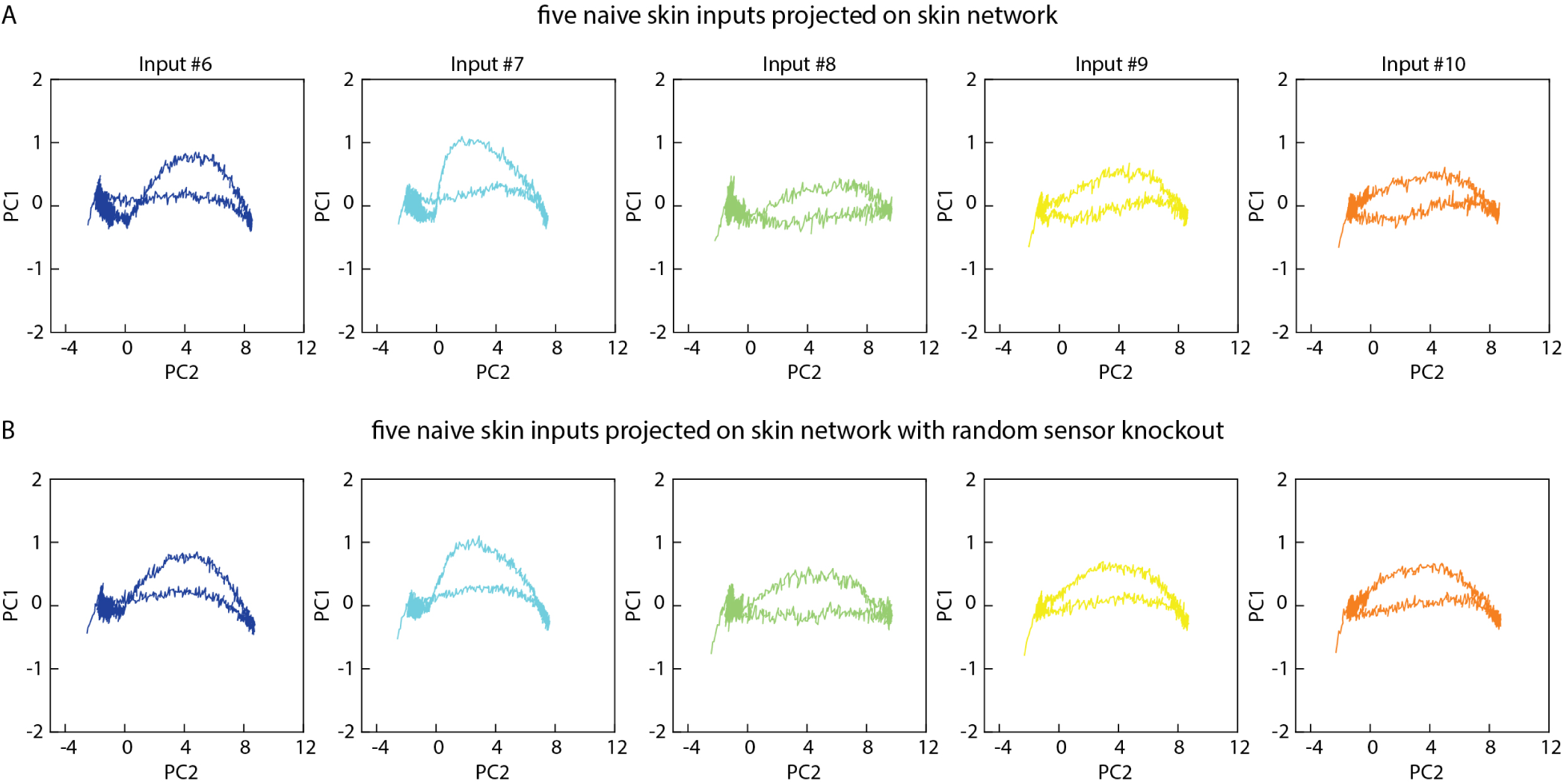
Effect of sensory knockout on the neuron population activity in the skin network. (**A**) The output of the neuron population for five different naive skin inputs (Fig. 1F-J) projected on the first two principal components (PCs). The plots illustrate outputs for the skin networks (same plots as in Fig. 3C, but here shown separately for each input rather than superimposed). (**B**) Similar display as in **A**, but with ‘knock-out’ of two random skin sensors post-learning.

**Supplementary Figure 7:**
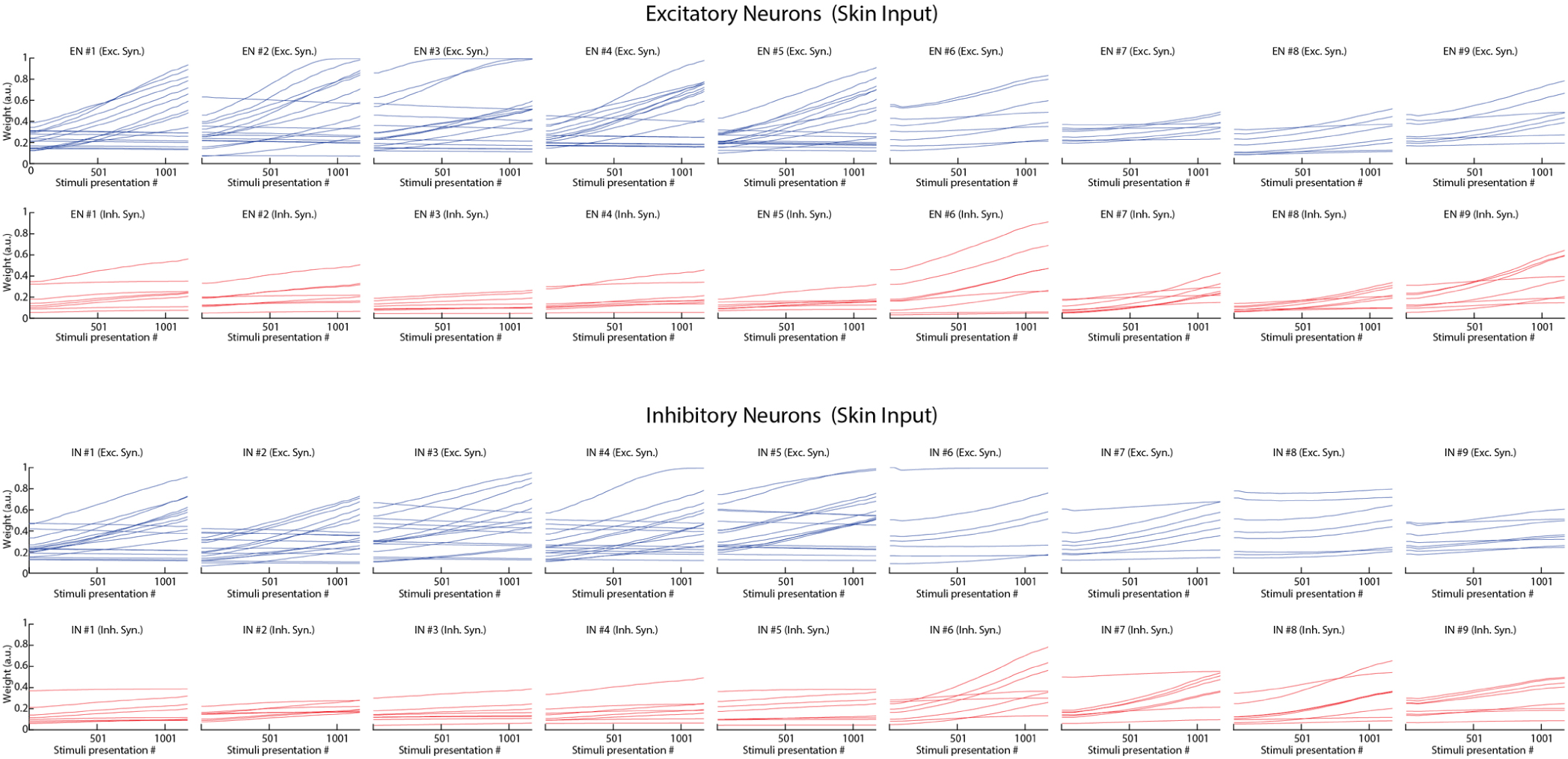
Excitatory and inhibitory synaptic weight evolution across all 18 neurons (9 EN & 9 IN) in the network, for learning based on skin inputs. The initial weights (weight distribution at stimulus presentation #1) correspond to the weights matrix in Fig. 4A. The end weights correspond to the weight matrix in Fig. 4B

**Supplementary Figure 8:**
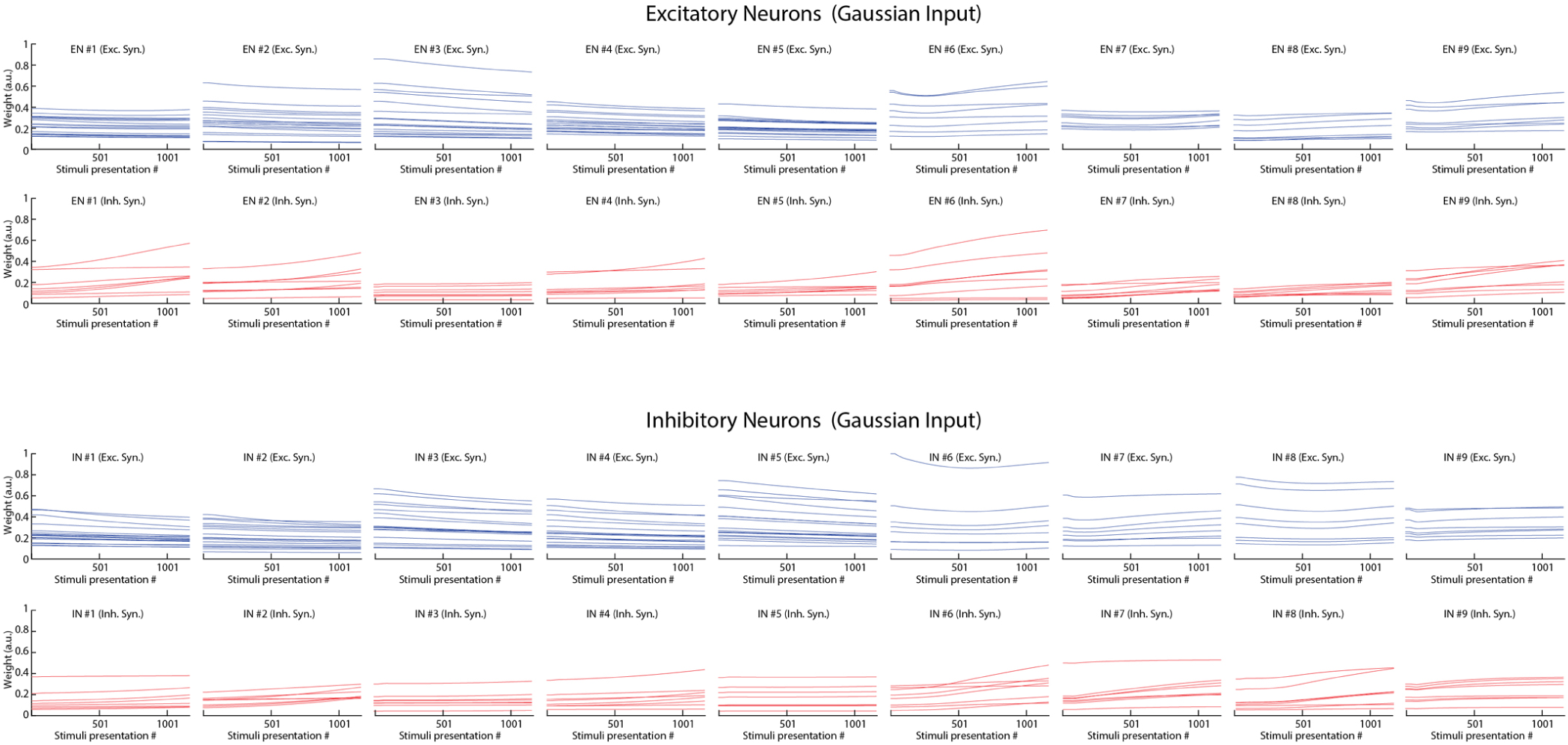
Excitatory and inhibitory synaptic weight evolution across all 18 neurons (9 ENs & 9 INs) in the network, for learning based on Gaussian inputs. The initial weights (weight distribution at stimulus presentation #1) correspond to the weights matrix in Fig. 4A. The end weights correspond to the weight matrix in Fig. 4C.

**Supplementary Figure 9:**
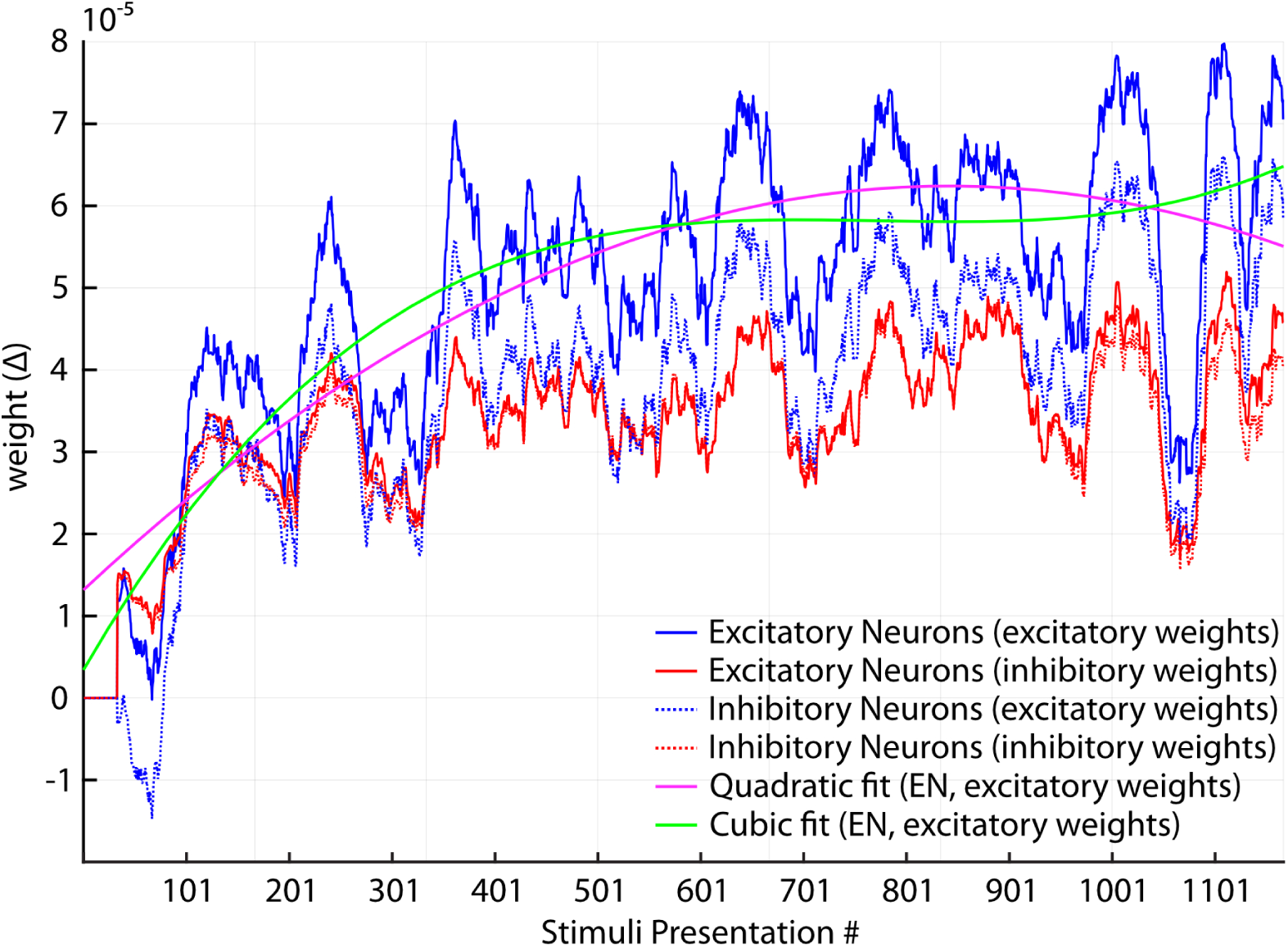
Excitatory and inhibitory synaptic weight change for all the stimuli presentation, across all 18 neurons (9 ENs & 9 INs) in the network.

**Supplementary Figure 10:**
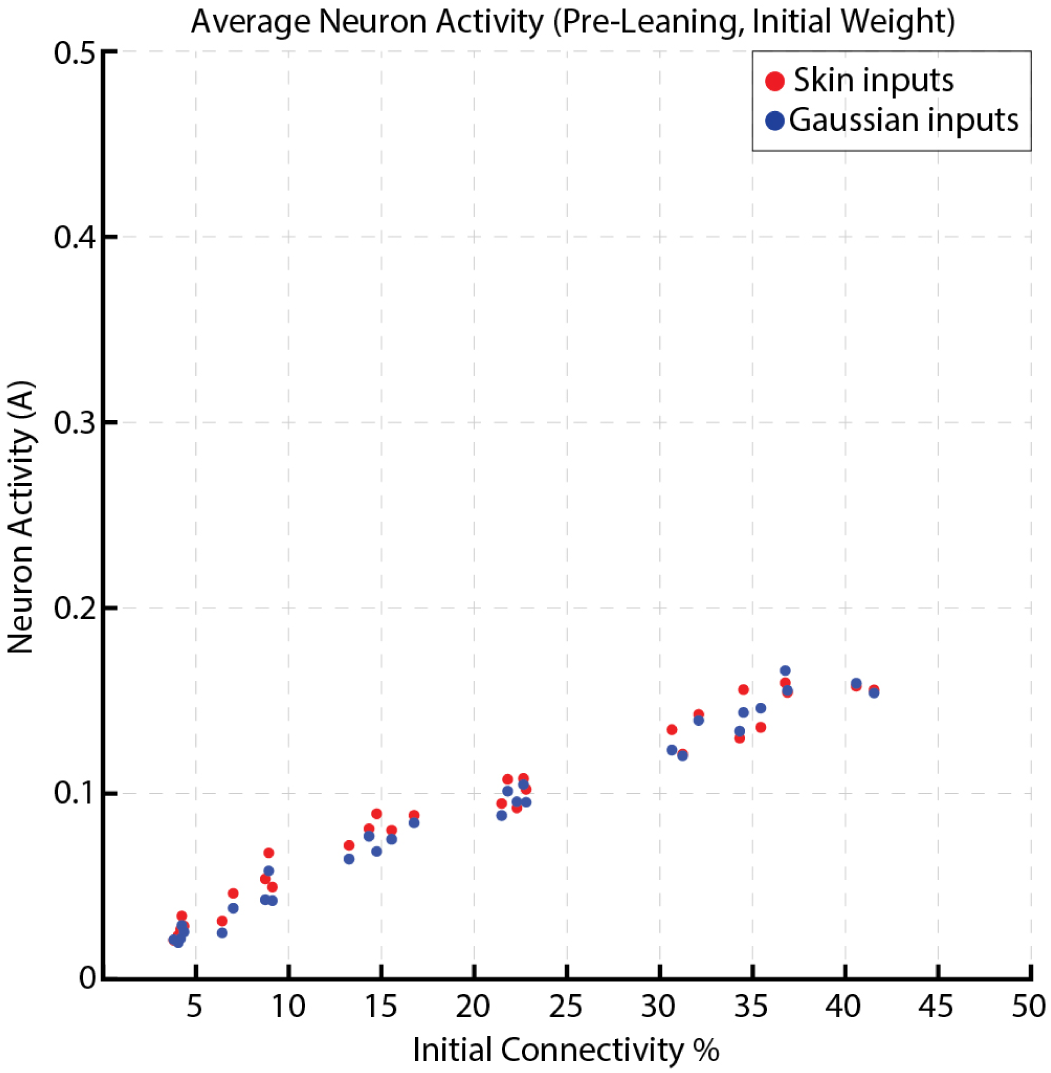
The average activity across all the neurons in the network for a given initial weight connectivity % and multiple sensory inputs (skin input & Gaussian input)

**Supplementary Figure 11:**
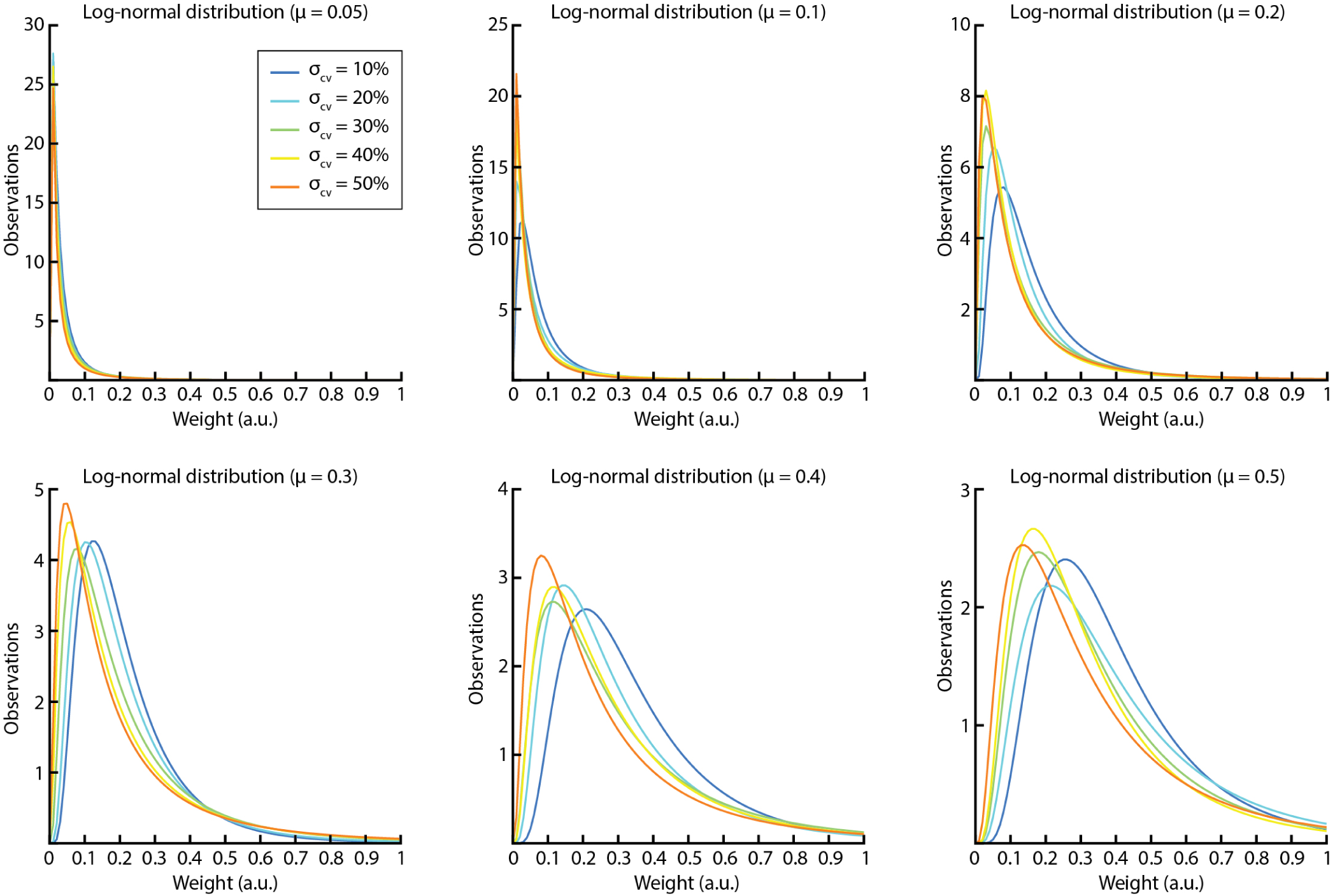
The 30 different initial weight settings tested. We used six different mean weights (*µ*; ranging between 0.05 & 0.5). For each mean weight, we used five different levels of synaptic weight spread, expressed as a the coefficient of variation (*σ_cv_*, ranging between 10% & 50%). The variance of the distribution of the initial weights was instead (*σ*) = (*σ_cv_/*100) ∗ *µ*. Each of the 30 different initial weight settings had its own randomization across the synapses, the numbers presented here only illustrates the data at the synaptic population level.

**Supplementary Figure 12:**
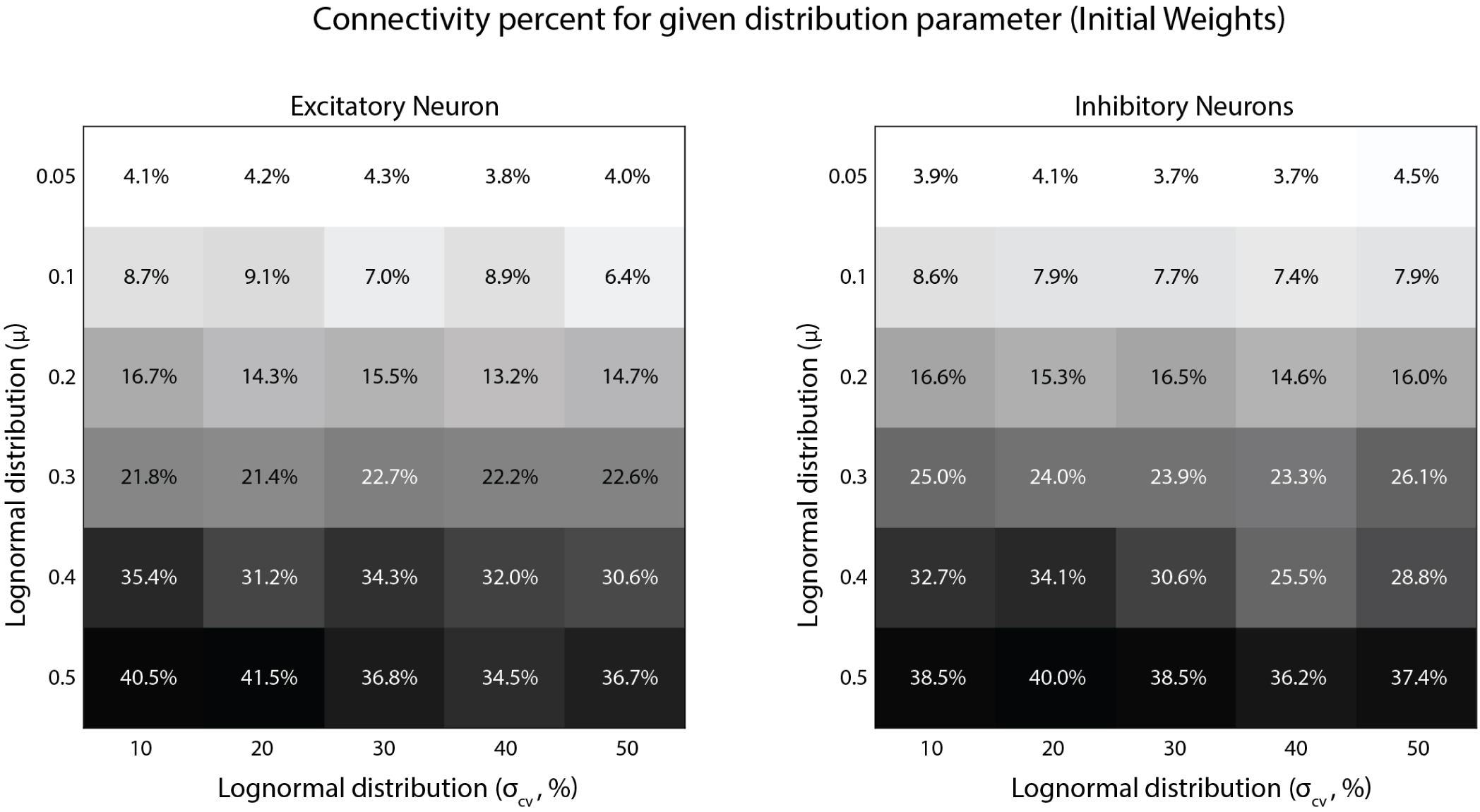
Network parameter settings for each specific Initial Connectivity% label. *µ* and *σ_cv_* refers to the distributions in Fig. S11. *Connectivity*% = Σ *initial weights / number of synapses*.

**Supplementary Figure 13:**
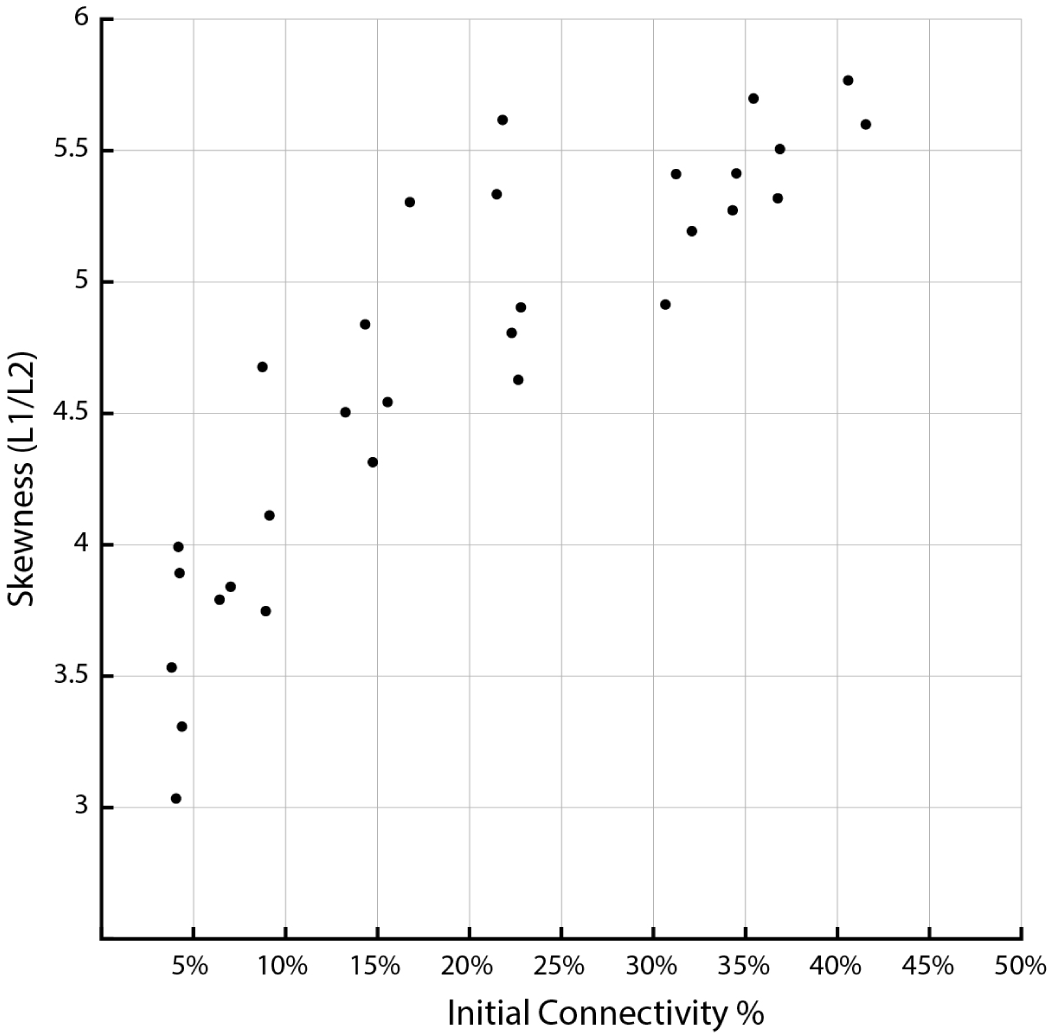
The skewness of initial synaptic weight distribution for a given connectivity %. The synaptic weight skewness was calculated as the ratio between the *l1* and *l2* norm of the weight vector (93). This measure will report its minimal value when all synapses have the same weight and a maximum value when one synapse has maximum weight and all other synapses have zero weight.

**Supplementary Figure 14:**
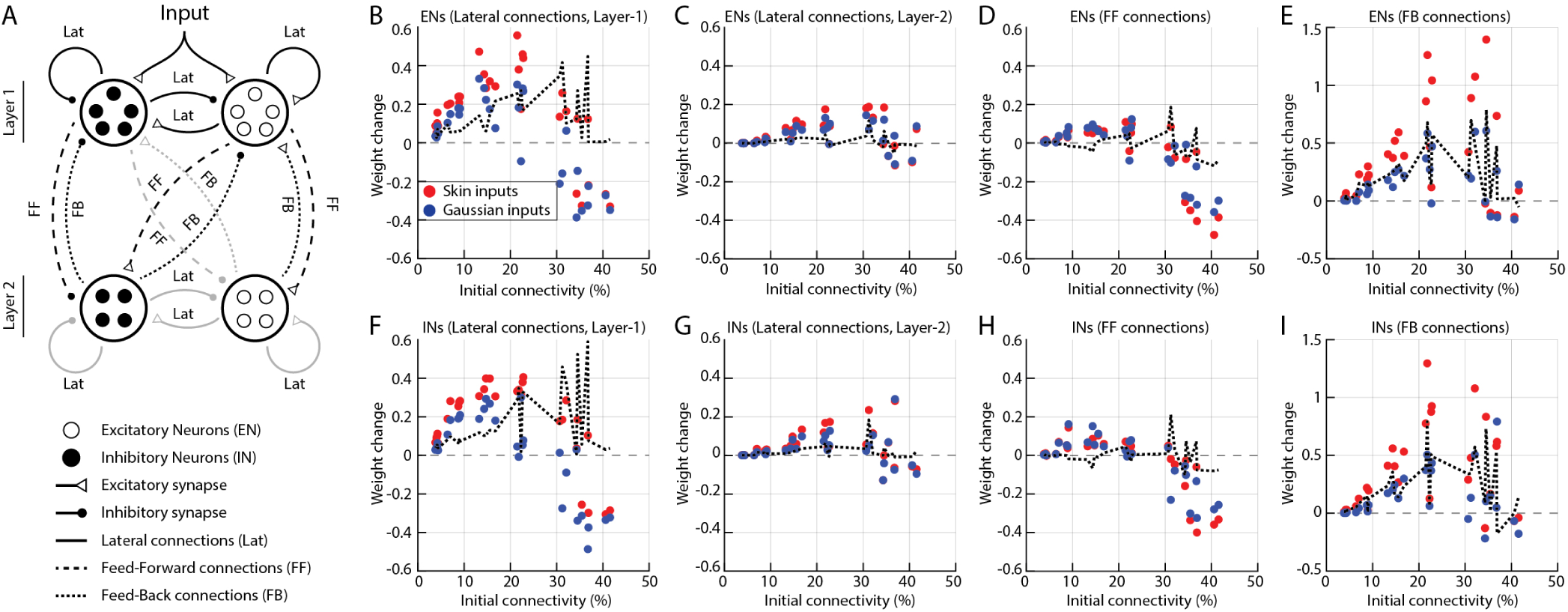
Synaptic weight changes resulting from learning for specific subtypes of synaptic connections. Same display conventions as in Fig. 4. (**A**) Schematic of the fully connected network structure studied; with specific labels for each type of connection. “*Lat*”: Lateral connections within a layer, “*FF*”: feed-forward connections from neurons in layer 1 to neurons in layer 2, “*FB*”: feed-back connections from neurons in layer 2 to neurons in layer 1. (**B**) Weight changes across all the excitatory and inhibitory lateral synaptic connections on excitatory neurons (*EN*) in layer 1. (**C**) Weight changes across all the lateral synapses on excitatory neurons in layer 2. (**D**) Weight changes across all the feed-forward synapses on excitatory neurons in layer 2. (**E**) Weight changes across all the feedback synapses on excitatory neurons in layer 1. (**F-I**) Weight change analysis similar to that in **B-E**, but instead for synaptic connections made on inhibitory neurons (*IN*). Overall, as in Fig. 4 the weights of all types of synaptic connections tended to increase, up to a certain level of mean initial connectivity (30%), above which the weights instead was decreased by the training. Across all conditions, except for the FF connections the weights increased more with the Skin input than with the Gaussian input (black dashed lines indicates the net difference between the two input conditions).

**Supplementary Figure 15:**
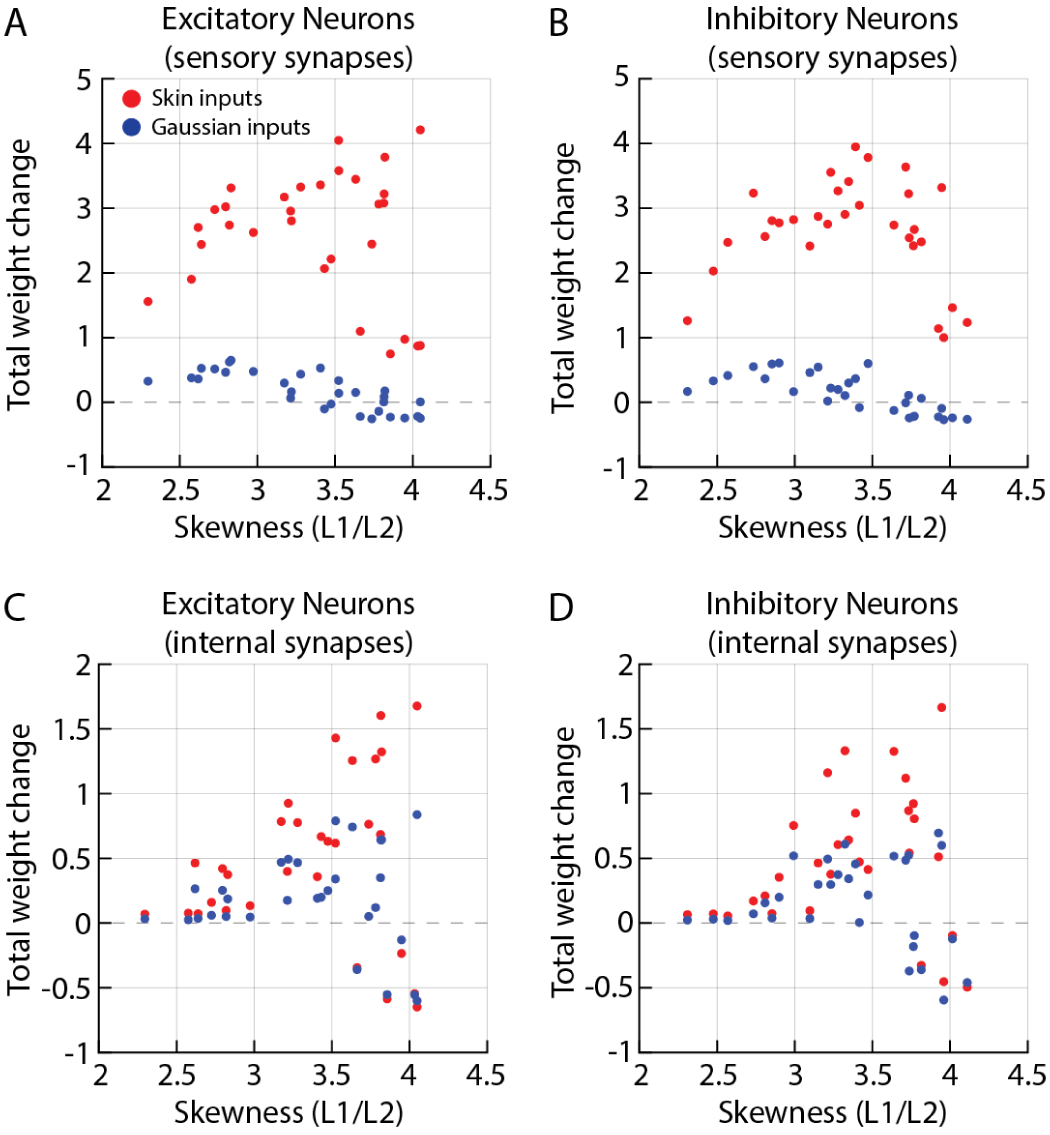
Effects of learning on synaptic weights. (**A**-**D**) The panels display average weight changes between pre-/postlearning, across all synapses of the indicated category (sensory synapses in **A**, **B** *and* internal synapses in **C**, **D**) for networks with varying skewness of initial synaptic weight distribution (Fig. S13) and sensory input type. Red markers indicate weight change within networks trained on skin inputs and blue markers for corresponding networks trained on Gaussian inputs. (**A**) Weight changes of the sensory synapses on ENs. (**B**) Weight changes of the sensory synapses on INs. (**C**) Weight changes of all excitatory and inhibitory internal synapses on ENs. (**D**) Weight changes of all excitatory and inhibitory internal synapses on INs.

**Supplementary Figure 16:**
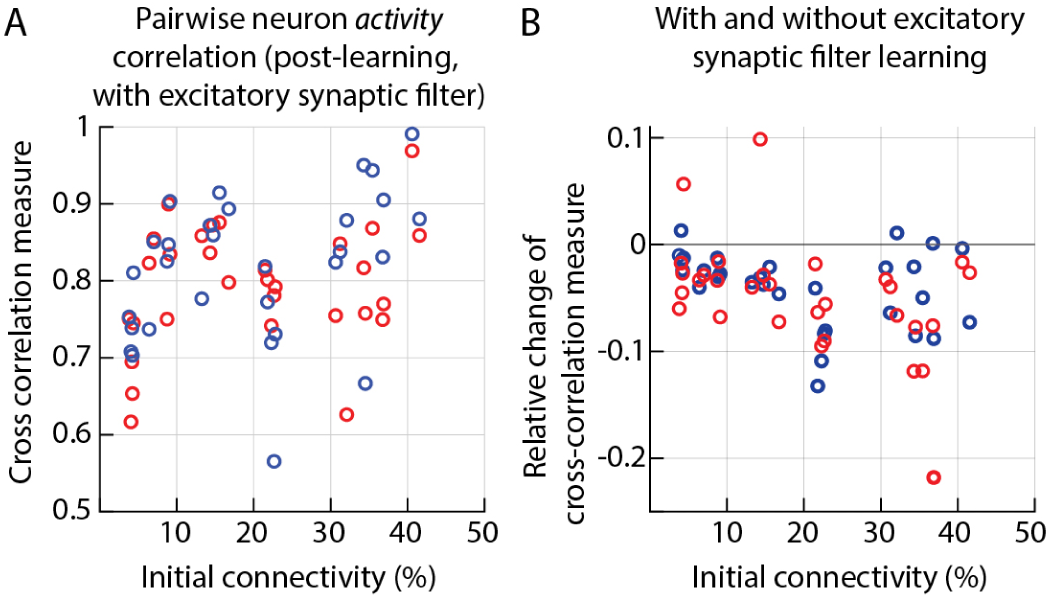
Effect of the excitatory synaptic learning signal filter on the neuronal cross-correlation. (**A**) Average pairwise cross-correlation measure across all the ENs activity in a network post-learning with excitatory synaptic learning signal filter of *e*^1.4^ (dotted vertical line in (Fig. 5C)), for networks with different initial connectivity and sensory inputs. (**B**) Net change in the cross-correlation measure of excitatory neuron activity caused by the learning signal filter (difference between (**A**) and Fig. 5B).

**Supplementary Figure 17:**
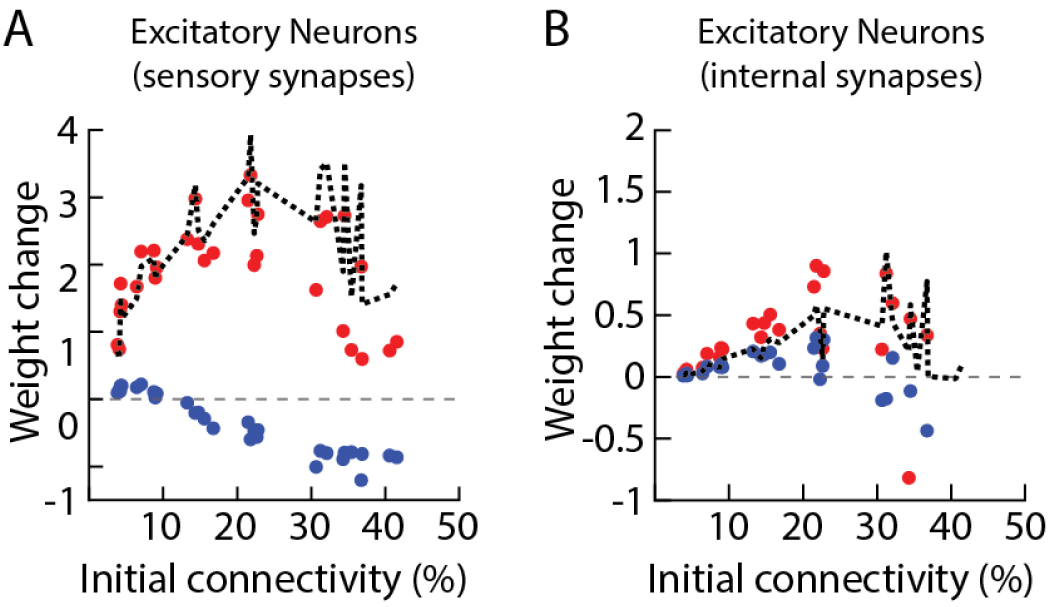
Effect of excitatory synaptic learning signal filter on synaptic weights. (**A**) Weight change across all the sensory synapses that project as excitatory synaptic connections on excitatory neurons in layer 1 for networks with different initial weight configurations. (**B**) Weight change across all the internal synapses that project as excitatory and inhibitory synaptic connections on all the excitatory neurons within the network. *Note:* **A & B** All networks were trained with the same excitatory synapse learning signal filter of *e*^1.4^ indicated by the vertical dotted line in Fig. 5B.

**Supplementary Figure 18:**
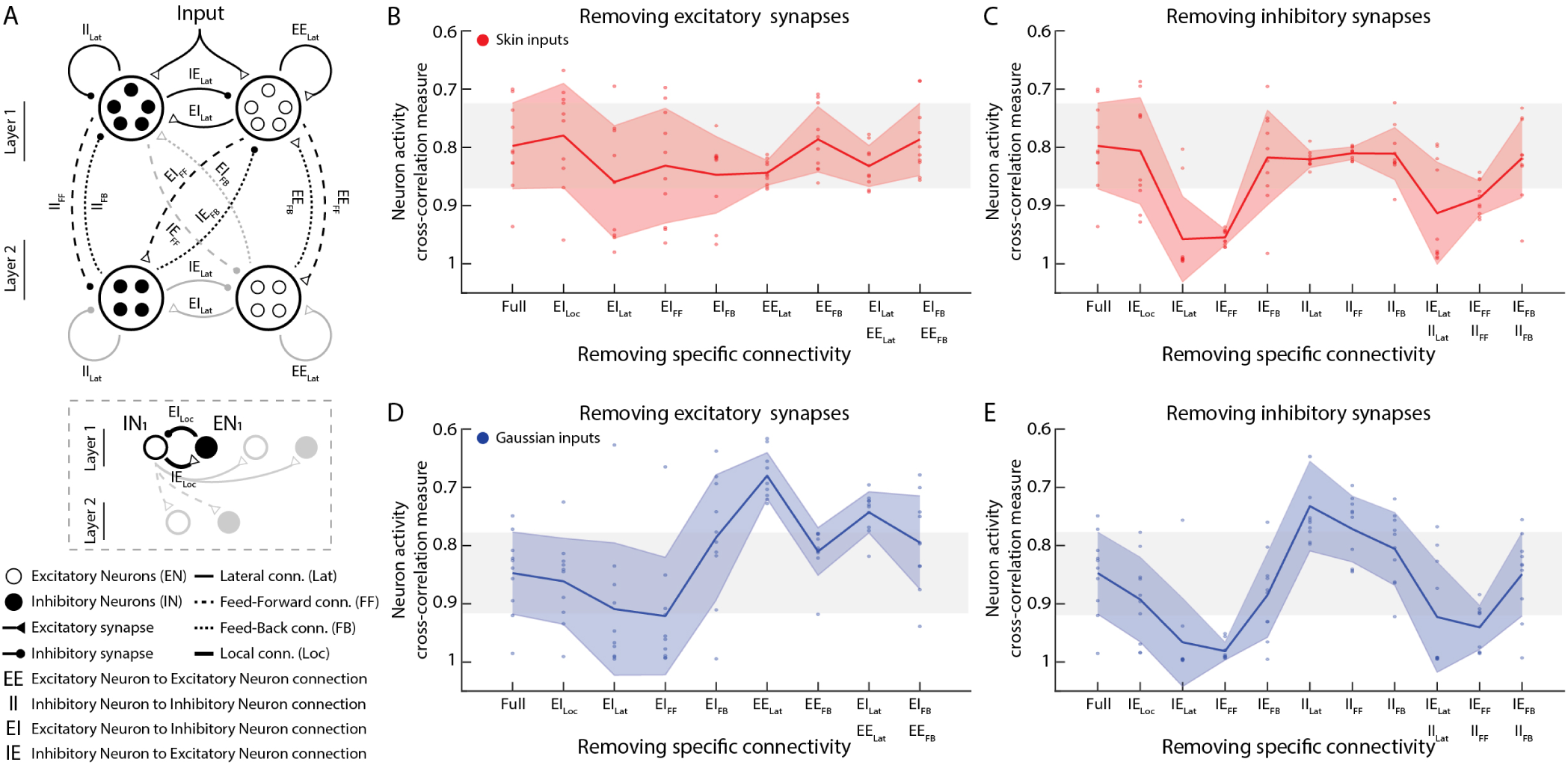
Impact of removing specific synaptic connections on learning quality. (**A**) Schematic of the fully connected network structure studied; with the insert showing specific labels for the type of individual excitatory and inhibitory synaptic connections between excitatory (ENs) and inhibitory neurons (INs). (**B**) The impact of removing specific types of outgoing excitatory neuron synaptic connections on the learning quality, as indicated by the average pairwise cross correlation measure post-learning. All learning is performed with skin inputs. *Note*: Full connectivity implies that the network has all the excitatory connections intact. The grey zone indicates the range of cross-correlation measures obtained with the Full connectivity. (**C**) The impact of removing specific types of outgoing inhibitory neuron synaptic connections on the learning. *Note*: Full connectivity implies that the network has all inhibitory connections. (**D, E**) Similar to **B, C**, where learning is performed with Gaussian inputs. The excitatory and inhibitory synaptic connections between ENs and INs were labelled as, “***EE***”: connections between excitatory - excitatory neurons, “***EI***”: connections between excitatory - inhibitory neurons, “***II***”: connections between inhibitory - inhibitory neurons, “***IE***”: connections between inhibitory - excitatory neurons. The topological connectivity between ENs and INs were labelled as, “***Lat***”: Lateral connections withing the ENs and INs in given layers, “***FF***”: feed-forward connections from ENs and INs in layer 1 to layer 2, “***FB***”: feed-back connections from ENs and INs in layer 2 to layer 1. (*insert*) “***Loc***”: Local connections between neighboring ENs and INs (local connection between EN*_i_* and IN*_i_*). The network has initial connectivity of 37% (log-normal distribution, *µ* = 0.5, *σ_cv_* = 10%, Fig. S11, S12). Ten different randomized initial weight vectors were generated for the same distribution parameters (individual marker for each given network configuration in **B-E**). All the network configuration learn with an excitatory synaptic filter of *e*^1.4^(Fig. 5A, dotted line).

**Supplementary Figure 19:**
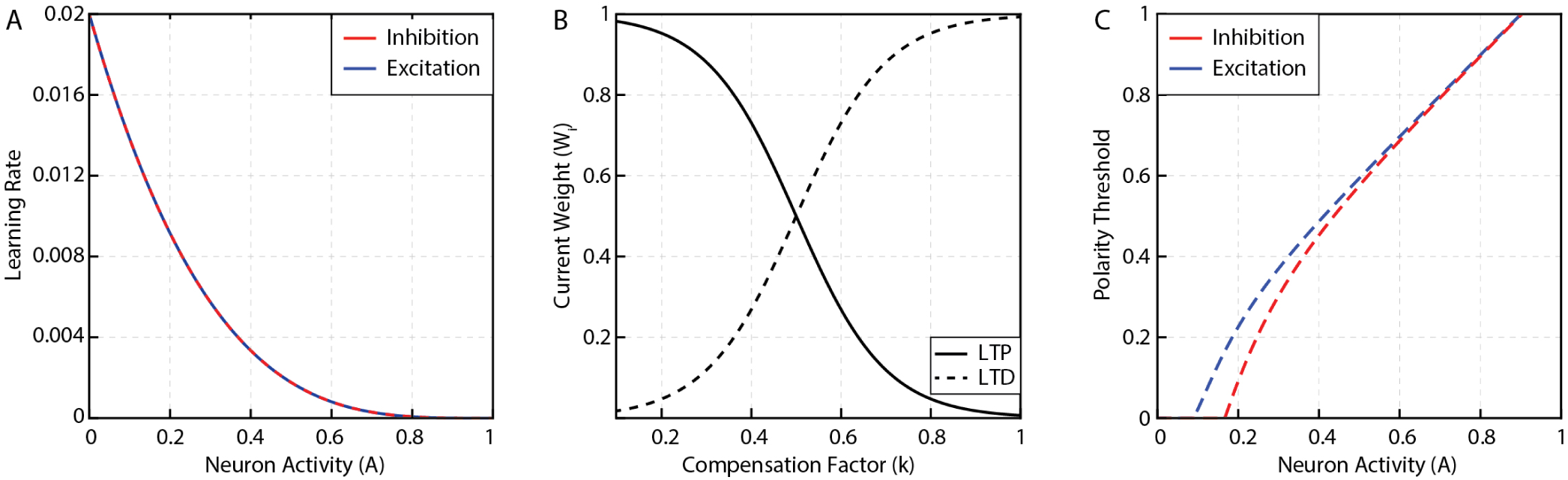
Illustration of multiple functions used for synaptic learning process in this study. (**A**) Learning rate as a function of neuron activity (*A*). (**B**) Compensation factor as function of current synaptic weight (*W_i_*). (**C**) Learning polarity threshold as a function of neuron activity (*A*).

